# Efficient stabilization of imprecise statistical inference through conditional belief updating

**DOI:** 10.1101/2022.06.08.495322

**Authors:** Julie Drevet, Jan Drugowitsch, Valentin Wyart

## Abstract

Statistical inference is the optimal process for forming and maintaining accurate beliefs about uncertain environments. However, human inference comes with costs due to its associated biases and limited precision. Indeed, biased or imprecise inference can trigger variable beliefs and unwarranted changes in behavior. Here, by studying decisions in a sequential categorization task based on noisy visual stimuli, we obtained converging evidence that humans reduce the variability of their beliefs by updating them only when the reliability of incoming sensory information is judged as sufficiently strong. Instead of integrating the evidence provided by all stimuli, participants actively discarded as much as a third of stimuli. This conditional belief updating strategy shows good test-retest reliability, correlates with perceptual confidence, and explains human behavior better than previously described strategies. This seemingly suboptimal strategy not only reduces the costs of imprecise computations, but counter-intuitively increases the accuracy of resulting decisions.

## Introduction

Efficient decision-making about the cause of noisy or ambiguous observations requires the accumulation of multiple pieces of evidence to form accurate beliefs [Wald, 1948; Bogacz, 2006], a process typically referred to as ‘statistical inference’. In stable environments, accumulating evidence across observations reduces uncertainty about their cause. There is ample experimental evidence that humans and other animals perform near-optimal inference in such conditions. However, human inference is subject to two main sources of internal noise: sensory noise which limits the amount of evidence provided by each observation [Green, 1966], and computation noise arising during the imprecise combination of incoming sensory evidence with the current belief [Drugowitsch, 2016; Wyart, 2016; Findling, 2021]. Both sources of noise trigger trial-to-trial variability in beliefs and behavior. In volatile environments, the cause of noisy observations changes over time, and statistical inference requires the appropriate weighting of the current belief against incoming sensory evidence [Glaze, 2015]. Recent findings suggest that humans and other animals are capable of performing near-optimal inference even in volatile environments [Murphy, 2021]. However, in these conditions, the trial-to-trial variability in beliefs triggered by internal sources of noise has even larger costs for efficient decision-making. Indeed, noisy sensory observations may be mistaken for genuine changes in their latent cause. Similarly, imprecise inference may trigger unwarranted changes-of-mind when incoming sensory evidence is combined imprecisely with internal beliefs. Whether and how humans may mitigate these significant cognitive costs in volatile environments remain unknown.

Here we addressed this question by studying human statistical inference in a volatile decisionmaking task based on noisy visual stimuli (Figure 1). Tested participants (*N* = 60 across two experiments) were presented with sequences of marbles which could be drawn either from a light bag containing dominantly light marbles or a dark bag containing dominantly dark marbles (Figure 1a). The light and dark areas of each marble were spatially scrambled, and the light/dark fractions of dominantly light and dark marbles were adjusted using an adaptive titration procedure (Figure 1b), such that 20% of dominantly light marbles (i.e., marbles from the light bag) were perceived as dark and vice versa (Figure 1c; see Methods). After each marble, participants were asked to identify the bag from which it was drawn (Figure 1d). Importantly, marbles were not drawn randomly and independently across successive trials, but rather in episodes of multiple draws from the same bag. Decision-making in the presence of such temporal structure can benefit from statistical inference that integrates uncertain visual information provided by each new marble with internal beliefs about the bag that is currently being drawn from.

**Figure 1.**
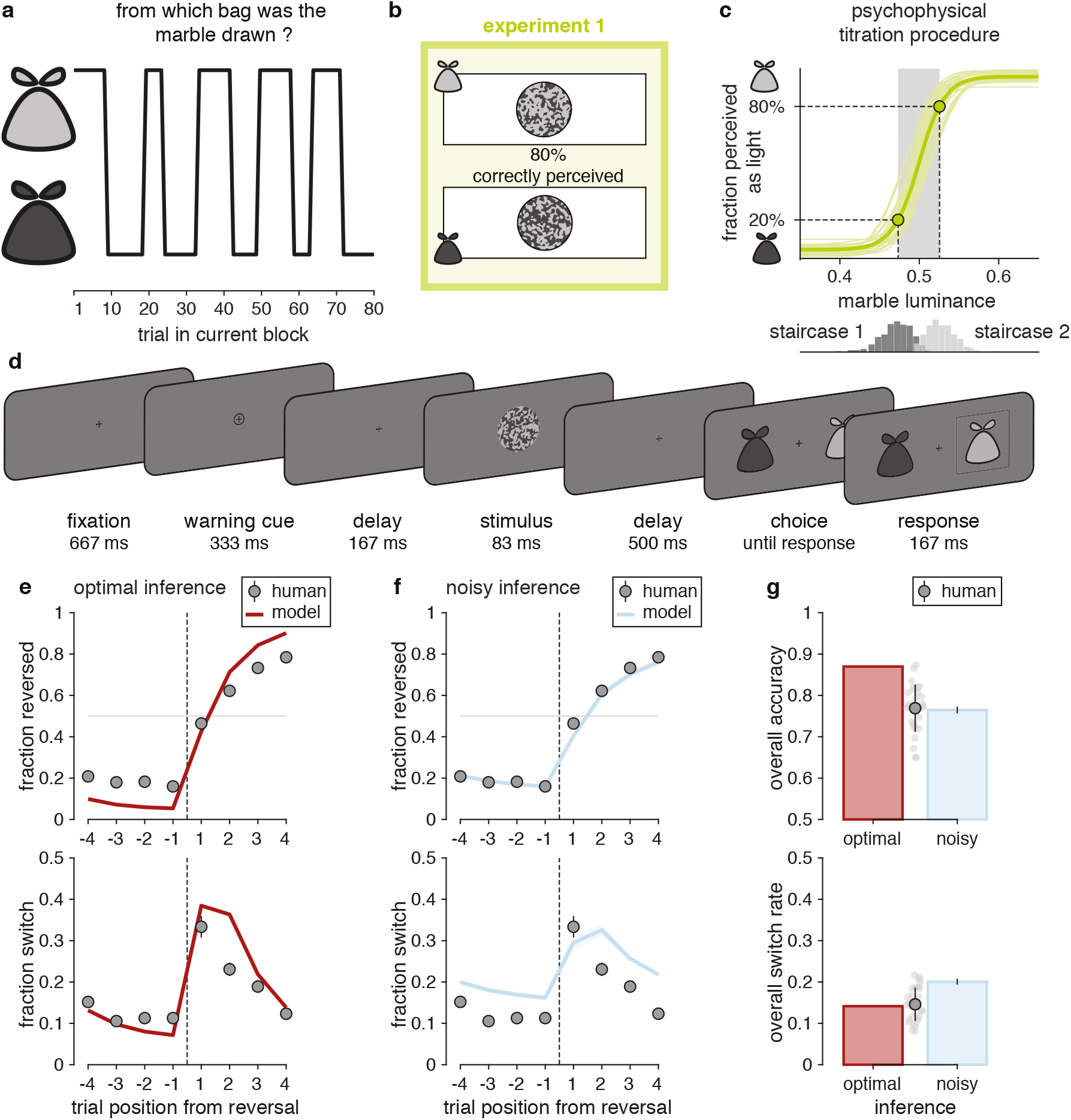
Description of experiment 1 (*N* = 30). **(a)** Structure of the volatile decision-making task. Each block of trials features alternations between draws from the light and dark bags. **(b)** Visual stimuli. The overall marble luminance is titrated such that 20% of presented marbles are miscategorized. **(c)** Adaptive titration procedure. Top: psychometric curves estimated by the titration procedure (thin lines: participants; thick line: group-level average). Bottom: distributions of marble luminance presented by the staircases titrating the dark and light marbles. **(d)** Trial description. 500 ms after a warning cue, a marble stimulus is presented for 83 ms, after which the participant indicates the bag from which marbles are currently drawn. **(e)** Predictions of optimal inference. Top: response reversal curves indicating the fraction of correct responses toward the new drawn bag surrounding each reversal. Bottom: response switch curves indicating the fraction of trial-to-trial response switches surrounding the same reversals. The reversal is represented by the thin dotted line. Dots indicate human data (group-level average), whereas lines indicate predictions of optimal inference. Error bars correspond to s.e.m. **(f)** Predictions of the noisy inference model. Top: response reversal curve predicted by the best-fitting noisy inference model (blue line). Bottom: response switch curve predicted by the best-fitting noisy inference model (blue line). Noisy inference captures well the accuracy of behavior surrounding reversals (top), but overestimates the variability of the same behavior (bottom). Same conventions as in (e). Shaded areas correspond to s.e.m. (g) Discrepancies between models and human behavior. Top: overall accuracy of participants (gray dot with error bar indicate the mean and s.d. of all participants, light gray dots indicate single participants), optimal inference (red bar) and noisy inference (blue bar). Bottom: overall switch rate of participants, optimal inference and noisy inference. Despite their suboptimal accuracy participants make response switches as often as optimal inference. Error bars on simulated noisy inference model (blue bars) correspond to s.e.m.

To analyze human behavior in this task, we developed a process-level model of statistical inference derived from normative Bayesian computations [Glaze, 2015], which hypothesizes distinct sources of internal noise at sensory, inference, and response selection stages [Drugowitsch, 2016]. Across two experiments, we fitted this model to human behavior to determine whether and how tested participants adopted strategies to compensate for the costs of imprecise statistical inference identified above. To do so, we formulated and compared different strategies by which participants could mitigate the variability of beliefs formed through statistical inference. We obtained converging evidence that human observers control statistical inference in a cost-efficient fashion, by updating their internal beliefs based on incoming inputs only when the noisy sensory observations are deemed sufficiently reliable. This metacognitive strategy not only reduces the costs of internal sources of variability affecting inference, but also counter-intuitively increases the accuracy of resulting decisions.

## Results

### Characterizing the suboptimality of statistical inference

We first assessed participants’ performance by comparing their reversal behavior to the one predicted by the optimal Bayesian inference process3 (see Methods, Eqs. 1 and 2). Importantly, this normative process accounts for sensory errors due to visual noise. As expected from previous work [Drugowitsch, 2016; Wyart, 2016], optimal inference substantially outperformed participants (Figure 1e,g; overall accuracy, optimal inference: 0.870; participants: 0.769 ± 0.011, mean ± s.e.m., paired two-sided *t*-test, difference: *t*_28_ = 9.5, *p* < 0.001, Cohen’s *d* = 1.858, 95% CI = [0.0790.123]). To characterize this suboptimality, we started from the optimal model but corrupted the inference with imprecise computations by adding normally-distributed noise on the result of each update step (Eq. 4). This suboptimal Bayesian inference model is controlled by two free parameters: 1. the hazard rate *h* – i.e., the subjective rate of reversals of the bag from which marbles are drawn, and 2. the inference noise *σ* – i.e., the standard deviation of these computation errors in the statistical inference process. On each trial, the model updates imprecisely its belief regarding the drawn bag by combining its prior belief in that trial with the imperfect information provided by the new visual stimulus – i.e., its likely bag based on its perceived lightness (see Methods).

We fitted this noisy inference model to two characteristic features of human reversal behavior simultaneously, each evaluated as a function of the time step from a change in hidden state (i.e. a reversal) on 4 trials preceding and 4 trials following each reversal (Figure 1f; see Methods for details): 1. the response reversal curve, corresponding to the fraction of correctly reversed responses regarding the new hidden state after reversal, and 2. the response switch curve, corresponding to the fraction of participants’ response switches regarding their own response on the previous trial. These two behavioral effects of interest are not independent, but each accounts for a separate behavioral dimension: the response reversal curve measures the accuracy of participants’ responses regarding the bag being drawn after a reversal, whereas the response switch curve measures the stability of participants’ responses across successive trials [Palminteri, 2017].

Simulations of the best-fitting noisy inference model provided a good fit to participants’ response reversal curves, but a poor fit to the same participants’ response switch curves when both features were fitted simultaneously. Indeed, the model showed the same overall accuracy as participants (Figure 1g; 0.765 ± 0.008, difference: *t*_28_ = 0.8, *p* = 0.416, Cohen’s *d* = 0.153, 95% CI = [−0.015 + 0.006], *BF*_01_ = 3.707), but larger overall switch rate (Figure 1g; 0.201 ±0.007; participants: 0.146± 0.007; difference: *t*_28_ = 10.0, *p* < 0.001, Cohen’s *d* = 1.858, 95% CI = [0.044 0.066]). Despite their suboptimal accuracy, participants showed a similar switch rate as optimal inference (Figure 1g; 0.141; difference: *t*_28_ = 0.6, *p* = 0.553, Cohen’s *d* = 0.111, 95% CI = [−0.020 + 0.011], *BF*_01_ = 4.294). This discrepancy between the humans’ and the noisy inference model’s overall switch rates suggests that participants deployed an additional strategy to compensate for the variability of beliefs due to imprecise inference.

Before assessing the nature of this ‘response stabilization’ strategy, we validated that participants’ suboptimal performance is due to imprecise inference. First, the hazard rate h inferred from the participants’ choices (0.081 ± 0.010) does not differ statistically from the true hazard rate of drawn bags (0.081; difference: *t*_28_ = 0.0, *p* = 0.999, Cohen’s *d* = 0.0002, 95% CI = [−0.020 +0.020], *BF*_01_ = 5.066). This suggests that participants don’t have a biased perception of the volatility of the task. Second, we ruled out noise in response selection as an alternative source of behavioral variability (Supplementary Figure 1a). Inference noise provides a better fit to behavior than selection noise (exceedance *p* > 0.999), a result which we validated through model recovery (see Methods). Third, we verified that participants’ suboptimal behavior was not best explained by a leaky integration process (Eq. 3) rather than by the normative one (Eq. 2). This linear approximation3 – even with additive inference noise – fails to better predict participants’ reversal behavior, and the suboptimal Bayesian model provides a better fit to both metrics as revealed by Bayesian model selection (Supplementary Figure 2a, exceedance *p* > 0.999).

### Comparing belief stabilization strategies for imprecise inference

We sought to characterize the cognitive strategy used by participants to make their responses more stable across successive trials. For this purpose, we derived five candidate strategies that implement such ‘stabilization’ at different processing stages in the noisy inference model described above (Figure 2), and compared them by simulating their specific effects surrounding reversals (Figure 3). The first candidate strategy we considered is a perceptual bias (Figure 2a), in which the sensory representation of each noisy stimulus is shifted in direction of prior beliefs following Bayes’ rule, resulting in a biased perception of the bag to which each stimulus belongs [Stocker, 2006; Luu, 2018; Lange, 2021] (see Methods, Eq. 5). This perceptual bias, operating prior to inference, leads to less variable beliefs at the expense of slower response reversals (Figure 3, middle column).

**Figure 2.**
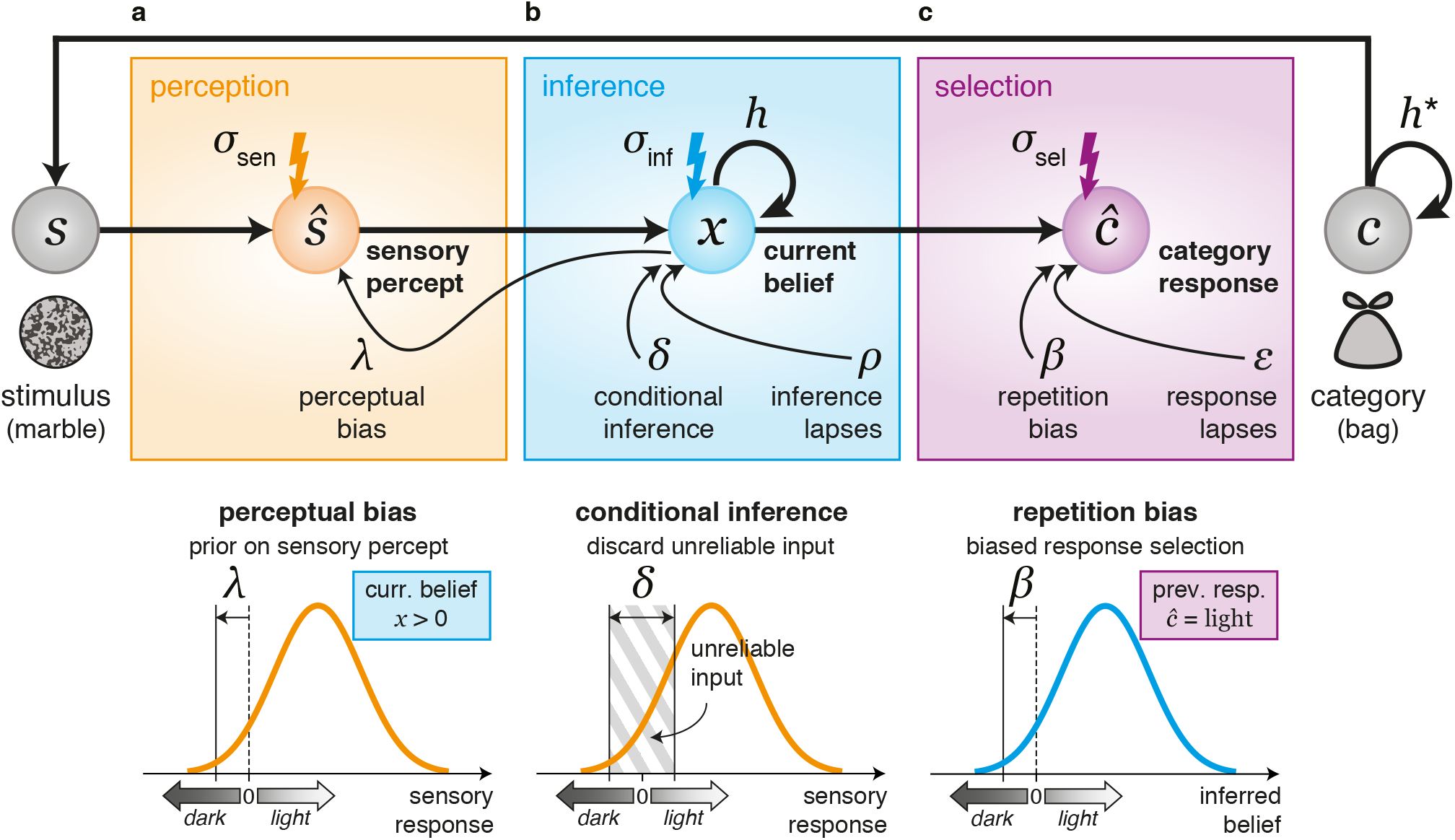
Candidate response stabilization strategies. A marble (stimulus *s*) is drawn from a given bag (category *c* = *lightordark*) at each trial. **(a)** At the perception stage, the continuous sensory response to the stimulus, corrupted by sensory noise *σ*_sen_, is categorized into a binary sensory percept *ŝ* (*light* or *dark*). A perceptual bias, controlled by parameter λ, biases the perceptual categorization toward the current belief *x* at stimulus onset by shifting the categorization criterion. **(b)** At the inference stage, the current belief *x* (expressed as the log-posterior odds ratio between the two bags) is updated as a function of the hazard rate *h* (prior term) and the incoming sensory percept *ŝ* (likelihood term). The inference process is corrupted by inference noise *σ*_inf_. Belief stabilization is achieved either through conditional inference, by discarding stimuli whose associated sensory responses do not exceed a reliability threshold *δ*, or through a random fraction ρ of random inference lapses during which the current belief is not updated. **(c)** At the selection stage, the current belief *x* is sampled with selection noise *σ*_sel_ to obtain a category response *ĉ*, corresponding to the bag perceived as being currently active (*light* or *dark*). Response stabilization is achieved either through a repetition bias toward the previous category response, controlled by parameter *β*, or through a random fraction *ϵ* of blind response repetitions (response lapses).

**Figure 3.**
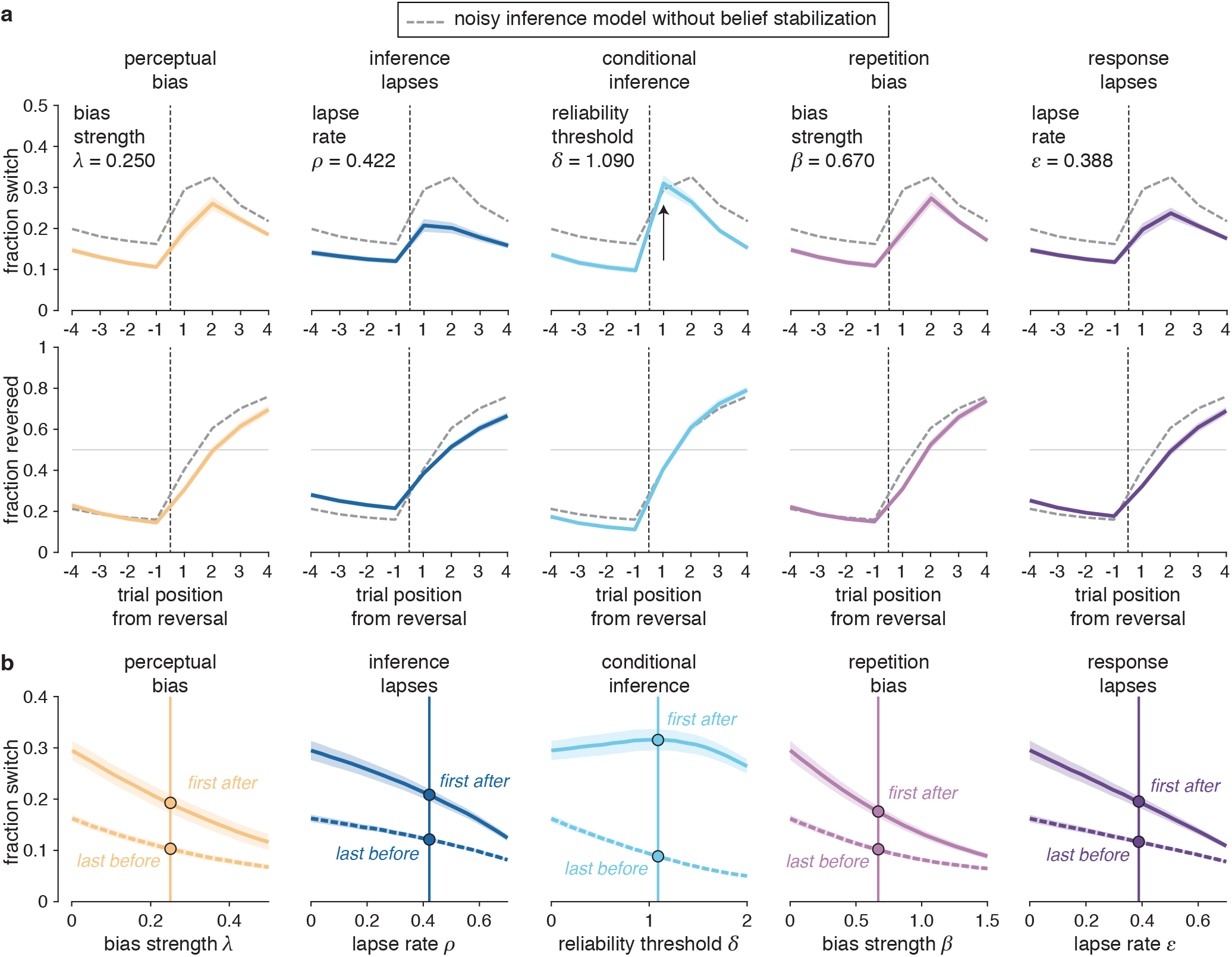
Predicted effects of response stabilization strategies. **(a)** Simulated effects of the five candidate strategies on response switch curves (top row) and response reversal curves (bottom row). The parameters controlling each response stabilization strategy (λ, *δ*, *ρ*, *β*, $*epsilon*) is set to match participants’ overall switch rate. The other parameters (*h*, *σ*_sen_, *σ*_inf_, *σ*_sel_) are fixed to their best-fitting values for the noisy inference model without response stabilization. Solid colored lines correspond to the reversal behavior of each candidate model, whereas dotted gray lines correspond to the reversal behavior of the noisy inference model without response stabilization (same for all panels). The conditional inference strategy stands out from other strategies on the first trial after reversal (arrow). **(b)** Simulated effects of the five candidate strategies on response switches just before and after a reversal. Fraction of response switches on the last trial before each reversal (dotted lines) and the first trial after each reversal (solid lines) for each response stabilization strategy. Dots correspond to stabilization parameters set as in (a). All candidate strategies reduce simultaneously response switches before and after reversals, except for conditional inference which reduces response switches only before reversals (i.e., when they are not warranted). Shaded areas around curves in (a) and (b) correspond to s.e.m.

The next two strategies we considered operate directly at the inference stage (Figure 2b). Instead of updating beliefs based on every noisy stimulus, we considered that participants may occasionally not update their beliefs, effectively ignoring the information provided by the presented stimulus and sticking to their prior beliefs (and their previous response) until the next trial. We contrasted two possible strategies resulting in such inference ‘omission’: 1. a fraction of inference lapses during which participants are distracted from the task (Eq. 6), or 2. a conditional inference strategy where participants update their beliefs only when the strength of the noisy sensory signal associated with the presented stimulus exceeds a certain threshold (see Methods, Eqs. 7 and 8). Although both strategies lead to less variable beliefs, making statistical inference contingent on the reliability of sensory representations does not delay response reversals by only discarding the stimuli that are more likely to be misperceived (Figure 3).

The last two considered strategies operate at the response selection stage, following the inference stage (Figure 2c): 1. a repetition bias which shifts participants’ response criterion in favor of the previous response (Eq. 9), and 2. a fraction of response lapses associated with ‘blind’ response repetitions (disconnected from current beliefs, Eq. 10). Like the perceptual bias and the inference lapses described above, these last two candidate strategies decrease the trial-to-trial variability of responses at the expense of slower adaptation to reversals (Figure 3).

Before identifying which of these candidate strategies best explains human behavior surrounding reversals (corresponding to the response reversal and switch curves described above), we assessed the ability to arbitrate between them through model recovery (see Methods). We generated synthetic data from each model and verified the ability to correctly recover the ‘ground-truth’ model through Bayesian model selection (Figure 4a; see Methods; all exceedance *p* > 0.95). We then compared the five response stabilization strategies in terms of their ability to fit response reversal and response switch curves: 1. quantitatively through Bayesian model selection (Figure 4b; see Methods), and 2. qualitatively by examining their best-fitting response reversal and switch curves (Figure 4c). Bayesian model selection revealed that the conditional inference strategy stands out by explaining simultaneously the accuracy (response reversal curve) and stability (response switch curve) of participants’ behavior surrounding reversals (Figure 4b; exceedance *p* > 0.999). Qualitatively speaking, while all five candidate strategies reproduce participants’ response reversal curves, only the conditional inference strategy explains the large and transient increase in response switches on the first trial following each reversal (Figure 4c). Besides, we confirmed that the alternative sub-optimal models (selection noise and leaky accumulation), even when stabilized with conditional inference, could not explain participants’ reversal behavior better than the noisy Bayesian inference model with conditional inference (Supplementary Figure 1b and Supplementary Figure 2b). These findings suggest that participants decrease the trial-to-trial variability of their beliefs by updating them only when the incoming sensory information is deemed sufficiently reliable – thereby discarding unreliable information that could otherwise trigger unwarranted changes-of-mind. In practice, the best-fitting reliability threshold *δ* (1.030 ± 0.122, mean ± s.e.m.) suggests that participants discard as many as 32.5% of presented stimuli as unreliable.

**Figure 4.**
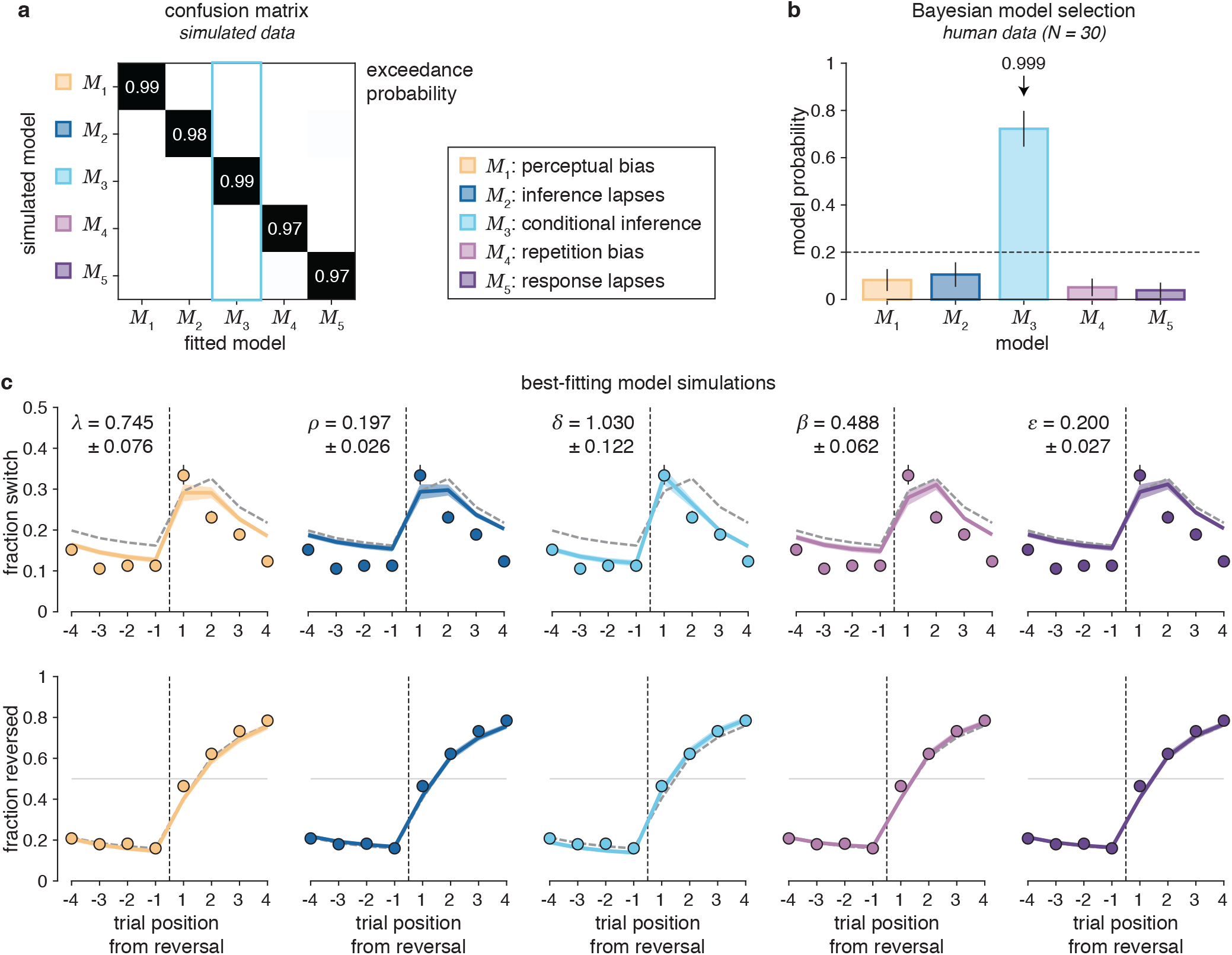
Belief stabilization through conditional inference (N = 30). **(a)** Confusion matrix between response stabilization strategies depicting exceedance probabilities pexc obtained from *ex-ante* model recovery. The ‘ground-truth’ response stabilization strategy used to simulate synthetic behavior was correctly recovered with *p*_exc_ > 0.97 for all five strategies, including conditional inference (the model that best describes participants’ behavior) with *p*_exc_ > 0.99. **(b)** Random-effects Bayesian model selection of the best-fitting strategy in participants’ data. Bars indicate the estimated model probabilities for the five candidate strategies. The conditional inference model (*M*_3_) best describes the behavior of more participants than other candidate strategies with *p*_exc_ > 0.999. Model probabilities are presented as mean and s.d. of the estimated Dirichlet distribution. The dashed line corresponds to the uniform distribution. **(c)** Simulations of response stabilization strategies fitted to participants’ data. Simulated response switch curves (top row) and response reversal curves (bottom row) based on the best-fitting parameters of each model. The mean best-fitting parameter (mean ± s.e.m.) is shown in the top-left corner for each subpanel. Solid colored lines correspond to the reversal behavior of each stabilized model, whereas dotted gray lines correspond to the reversal behavior of the noisy inference model without response stabilization (same for all panels). Dots indicate human data (group-level average). Only the conditional inference model reproduces participants’ reversal behavior. Shaded areas and error bars correspond to s.e.m.

### Validating specific predictions of conditional inference

To provide further evidence in favor of conditional inference, we designed and ran a second experiment (*N* = 30 new participants) which allowed: 1. replicating the findings obtained in experiment 1, and 2. validating two specific predictions of conditional inference. First, we reasoned that if participants discard less reliable stimulus information, they should do so more often when the stimulus is difficult to categorize as light or dark – and hence triggers a weaker sensory signal. To test this first prediction, we had each bag contain marbles of three levels of difficulty (stimulus strength, corresponding to different proportions of light and dark areas) determined through the same titration procedure to achieve correct stimulus categorization of 70%, 80% and 90% (Figure 5a). We predicted that participants should discard more difficult stimuli more often than easier stimuli.

**Figure 5.**
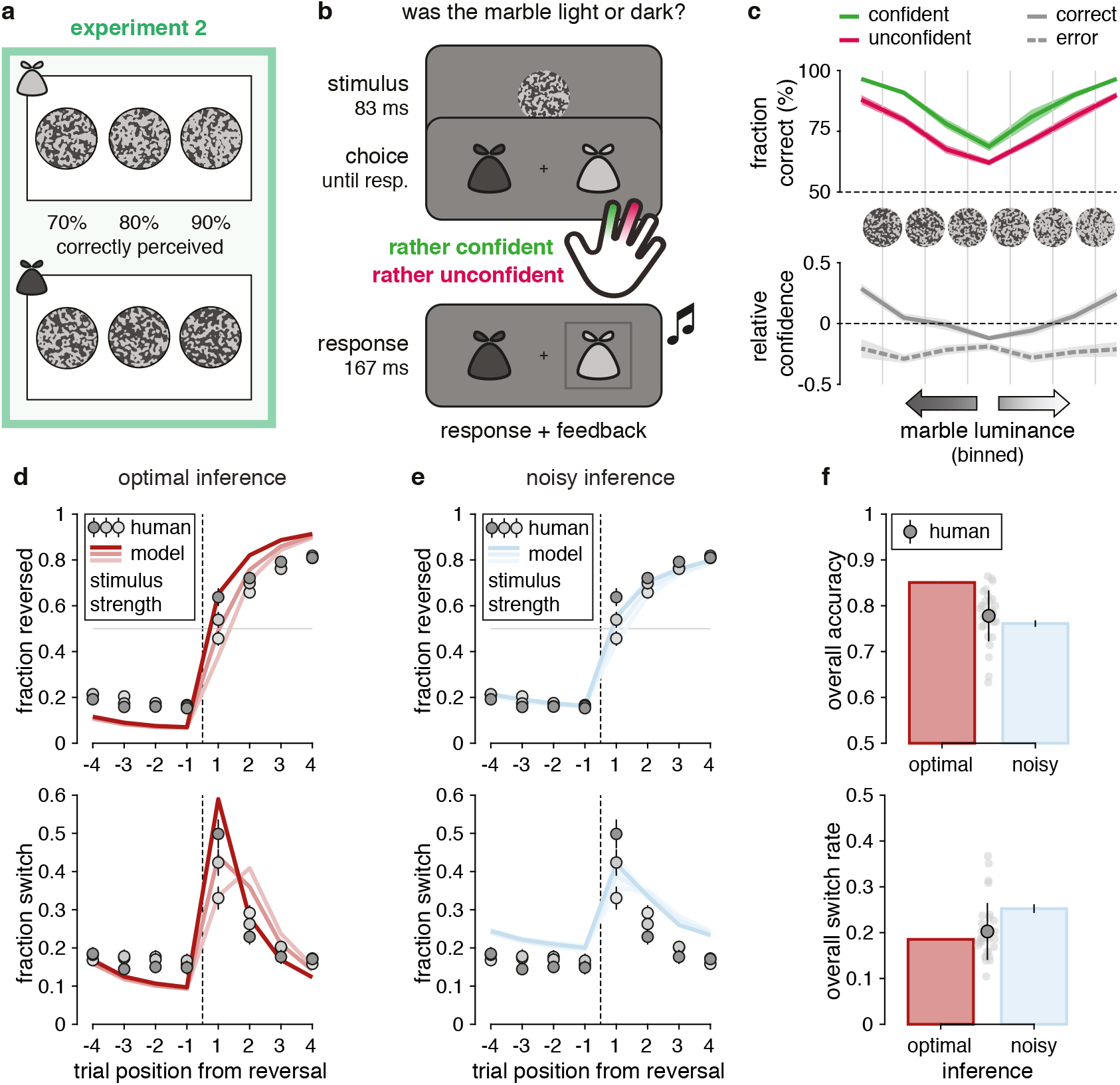
Description of experiment 2 (N = 30). **(a)** Visual stimuli. Each bag is filled with three types of marbles, corresponding respectively to 70%, 80% and 90% of correctly categorized stimuli, using the same adaptive titration procedure as in experiment 1. **(b)** Titration trials with confidence reports. Participants were asked to categorize the presented marble as light or dark, and report simultaneously their confidence level as rather high or low. Participants received auditory feedback after each titration trial. **(c)** Relation between decision confidence and accuracy in titration trials. For each binned luminance, participants’ fraction of correct decisions for confident (green) and unconfident (red) trials (top panel), and participants’ fraction of confident decisions for correct (solid line) and error (dashed line) trials (bottom panel). Confidence is expressed relative to each participant’s mean confidence (dashed line). Shaded areas correspond to s.e.m. **(d)** Predictions of optimal inference as a function of stimulus strength on the first trial following each reversal. Top: response reversal curves indicating the fraction of responses toward the new drawn bag surrounding each reversal. Bottom: response switch curves indicating the fraction of trial-to-trial response switches surrounding the same reversals. The reversal is represented by the thin dotted line. Dots indicate human data (group-level average), whereas lines indicate predictions of optimal inference. Error bars correspond to s.e.m. **(e)** Predictions of the noisy inference model as a function of stimulus strength on the first trial following each reversal. Noisy inference captures well the accuracy of behavior surrounding reversals (top), but overestimates the variability of behavior (bottom). Same conventions as in (d). Shaded areas correspond to s.e.m. **(f)** Discrepancies between models and human behavior. Top: overall accuracy of participants (gray dot with error bar indicate the mean and s.d. of all participants, light gray dots indicate single participants), optimal inference (red bar) and noisy inference (blue bar). Bottom: overall switch rate of participants, optimal inference and noisy inference. Despite their suboptimal accuracy, participants make response switches as often as optimal inference. Error bars on simulated noisy inference model (blue bars) correspond to s.e.m.

Second, we hypothesized that the reliability threshold used by participants to discard sensory information should decrease with participants’ confidence at categorizing isolated stimuli as light or dark. In other words, more confident participants should discard fewer stimuli as unreliable. For this purpose, we asked participants to provide confidence reports when categorizing stimuli during the titration procedure (Figure 5b; see Methods). These confidence reports showed classical signatures of decision evaluation (Figure 5c): lower accuracy for decisions made with low confidence (repeated-measures ANOVA, *F*_1,29_ = 84.9, *p* < 0.001, 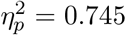), and a selective increase in confidence as a function of stimulus strength for correct decisions but not errors (correct:*F*_3,87_ = 61.8, *p* < 0.001, *η*^2^ = 0.681; error: *F*_3,87_ = 1.5, *p* = 0.241, *η*^2^ = 0.048).

As in experiment 1, participants showed suboptimal performance (Figure 5d) and imprecise inference overestimated the trial-to-trial variability of participants’ responses (Figure 5e), which was again closer to the overall behavioral variability of optimal inference (Figure 5f). In contrast to experiment 1, we could split response reversal and response switch curves as a function of stimulus strength on the first trial following each reversal. Model simulations showed that conditional inference increases the degree of separation between these curves (Supplementary Figure 3). This is because the growing fraction of discarded stimuli with categorization difficulty results in missed reversals when they are followed by a difficult stimulus. Model recovery confirmed that the conditional inference model is in principle identifiable from behavior (Supplementary Figure 4a; conditional inference *M*_3_: exceedance *p* > 0.99), and Bayesian model selection yielded strong evidence in favor of conditional inference (Supplementary Figure 4b; exceedance *p* > 0.999). As in experiment 1, conditional inference is the only strategy that can explain the dynamics of participants’ response switches surrounding reversals (Supplementary Figure 3 and Supplementary Figure 4c).

To validate our first prediction that participants discard more difficult stimuli more often than easier ones, we fitted a separate reliability threshold for each stimulus strength. The reliability threshold is expressed in abstract units, and a more tangible metric of conditional inference corresponds to the associated ‘discard rate’: the overall fraction of discarded stimuli predicted by the model (see Methods). The proposed conditional inference strategy predicts that the reliability threshold should not vary with stimulus strength (which varies from one trial to the next). We found that the reliability threshold remains indeed approximately constant (Figure 6a; *F*_2,58_ = 2.6, *p* = 0.095, *η*^2^ = 0.081), resulting in a decreasing discard rate with stimulus strength – from 33.3±3.8% (mean ± s.e.m.) for difficult stimuli categorized with 70% accuracy down to 13.6± 1.2% for easy stimuli categorized with 90% accuracy (*F*_2,58_ = 14.6, *p* < 0.001, *η*^2^ = 0.335). This finding supports the notion that participants control inference by ignoring the sensory information below a fixed reliability threshold.

**Figure 6.**
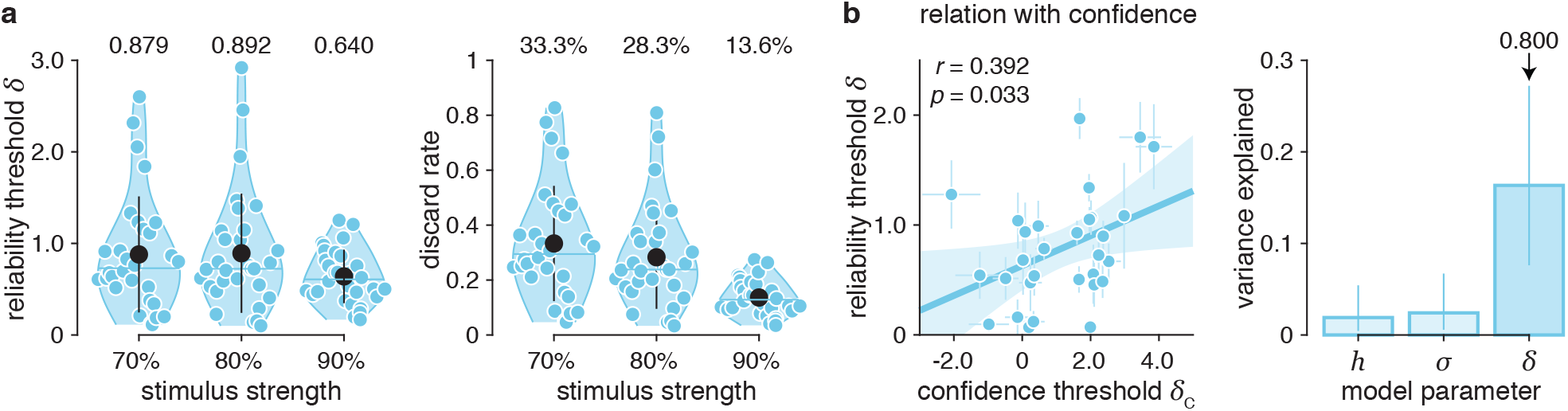
Validation of specific predictions of conditional inference (N = 30). **(a)** Decrease in evidence discard rate with stimulus strength. The reliability threshold does not vary with stimulus strength, resulting in discard rates which decrease with stimulus strength (one blue dot per participant). Black dots represent group-level means and error bars represent their associated standard deviations. Horizontal blue lines represent group-level medians. **(b)** Selective relation between reliability and confidence criteria. Left panel: correlation between confidence threshold and reliability threshold across participants. As predicted by conditional inference, the reliability threshold estimated in the volatile decision-making task correlates positively with the confidence threshold estimated when categorizing isolated stimuli. Parameters are presented as mean ± s.d. of posterior distributions of each fit. Shaded area corresponds to the 95% confidence interval for the regression line. Right panel: variance of confidence threshold explained by each model parameter. The reliability threshold shares more variance with the confidence threshold than the other two model parameters. Bars correspond to *r*^2^ and error bars to the interquartile ranges of each *r*^2^ measure obtained through bootstrapping (*N* = 10^4^).

To validate our second prediction that the reliability threshold used to discard unreliable sensory information should decrease with confidence across participants, we fitted the confidence threshold *δ_c_* of each participant during the titration procedure (Supplementary Figure 5; see Methods). We then regressed this confidence parameter – which decreases with confidence – against the parameters of the conditional inference model fitted to the same participants during the main task. As predicted, we found a positive relation between the confidence threshold and the reliability threshold across participants (Figure 6b; linear correlation coefficient *r* = 0.392, *p* = 0.033, 95% CI = [0.037 0.659]). This positive relation remained significant when accounting for the apparent internal variability in confidence reports (Supplementary Figure 5c; *b* = 0.250±0.089, *p* = 0.009, 95% CI = [0.0750.425]). This second finding suggests that the reliability threshold used to discard unreliable sensory information depends on the confidence with which participants categorize individual stimuli.

### Identifying individual differences in conditional inference

To provide further support for conditional inference, we studied whether the reliability threshold used to discard stimuli shows specific individual differences across participants. We reasoned that, if individual differences in reliability threshold *δ* reflects genuine differences in conditional inference, then these differences should follow a behavioral ‘gradient’ that is distinct from the ones associated with the other two parameters in the model (the hazard rate *h* and the inference noise *σ*). More specifically, individual differences in *δ* should mainly affect the variability of participants’ responses surrounding reversals – i.e., their response switch curves.

First, we observed that all three parameters show large individual differences (Figure 7a). Importantly, these parameters are only moderately correlated with each other (Figure 7b) – with at most 10% of shared pairwise variance – and each of them shows significant test-retest reliability across the two experimental sessions performed by each participant on separate days (hazard rate *h*: *r* = 0.698, *p* < 0.001, 95% CI = [0.5380.809]; inference noise *σ*: *r* = 0.308, *p* = 0.018, 95% CI = [0.0560.523]; reliability threshold *δ*: *r* = 0.574, *p* < 0.001, 95% CI = [0.3730.724]; Supplementary Figure 6; see Methods). To test whether each parameter is associated with a specific behavioral gradient, we split participants in two groups as a function of the best-fitting value of the parameter being considered and plotted the response reversal and switch curves for each group (Figure 7c; see Methods). We found that the behavioral gradient associated with the hazard rate *h* explains 48.7% of individual differences in response reversal curves, with a clear effect on the reversal time constant. The behavioral gradient associated with inference noise *σ* explains 22.5% of individual differences in response reversal curves, with a distinct effect on the overall accuracy of responses. By contrast, the behavioral gradient associated with the reliability threshold *δ* explains only 4.2% of individual differences in response reversal curves, but 24.8% of individual differences in response switch curves. In line with the conditional inference model, participants with higher reliability thresholds show less variable responses but equally rapid adaptation to reversals (Figure 7c).

**Figure 7.**
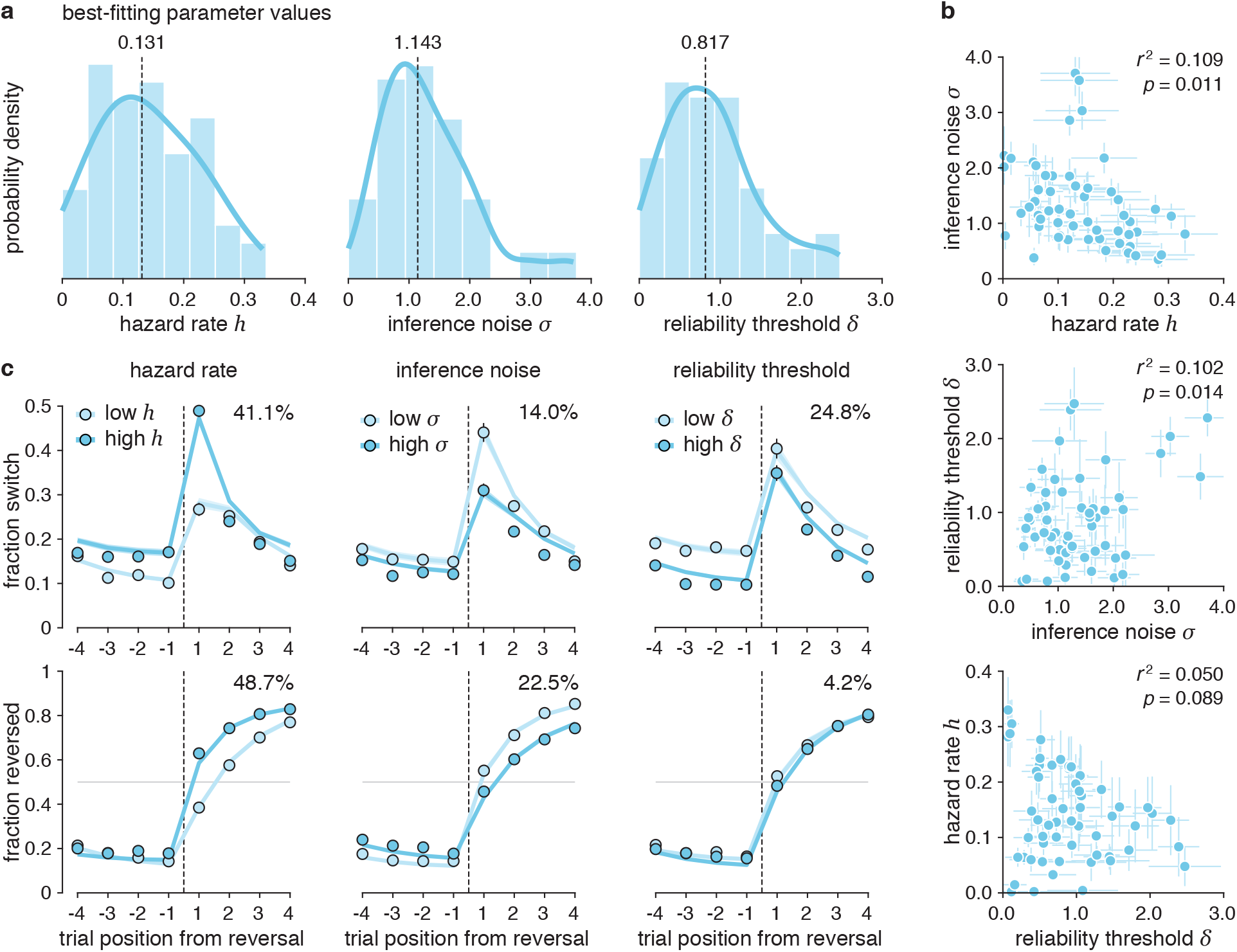
Interindividual variability in conditional inference (N = 60). **(a)** Distribution of conditional inference model parameter values across participants from both experiments. Best fitting parameters and their median value (dotted lines) with density approximation (solid blue lines). **(b)** Pairwise relations between the parameters of the conditional inference model. Parameters are only mildly correlated with one another. Parameters are presented as mean ± s.d. of posterior distributions of each fit **(c)** Distinct gradients of behavioral variability associated with each parameter of the conditional inference model (left: hazard rate; middle: inference noise; right: reliability threshold). Response switch curves (top) and response reversal curves (bottom) obtained by sorting participants in two median-split groups as a function of the best-fitting value of each parameter. Lines correspond to simulations of the best-fitting conditional inference model, whereas dots correspond to participants’ behavior. Percentages indicate the variance in response switch curves (top row) and response reversal curves (bottom row) explained by each parameter. Shaded areas and error bars correspond to s.e.m.

To validate the existence of this behavioral gradient in a ‘bottom-up’ fashion – independently of fits of the conditional inference model, we performed a principal component analysis (PCA) of participants’ response reversal and switch curves (Supplementary Figure 7; see Methods). Strikingly, the three first principal components (PC) obtained using this variance partitioning procedure map on the three parameters of the conditional inference model. The third PC (PC3) is associated with individual differences in response switch curves (16.4%), but little to no variability in response reversal curves (3.4%). Accordingly, participant-specific scores on this PC correlate selectively with reliability thresholds fitted with the conditional inference model (*r*^2^ = 0.348, exceedance *p* = 0.861, 95% CI = [0.1680.551]). Together, these top-down (model-based) and bottom-up (PCA-based) findings provide additional evidence for conditional inference by identifying a behavioral gradient characteristic of conditional belief updating across tested participants.

### Measuring cognitive benefits of conditional inference

The reduced trial-to-trial variability of beliefs triggered by conditional inference mitigates the negative effects of external (unreliable sensory information) and internal (imprecise computations) sources of variability during statistical inference, by discarding the incoming stimulus information that does not exceed a minimum reliability level. We thus tested whether conditional inference may not only decrease the variability of beliefs and resulting responses, but also counter-intuitively increases accuracy. For this purpose, we compared the effects of the five tested belief stabilization strategies on accuracy across both experiments through simulations (Figure 8a). We found that all strategies decrease behavioral variability at the expense of lower accuracy, except for conditional inference which shows an inverted U-shaped relation with accuracy. The accuracy-optimizing value *δ*\* of the reliability threshold (*δ*\* = 1.281, associated with 5.5% more accurate responses) is associated with as much as 41.3% of discarded stimuli categorized with an overall 80% accuracy (Supplementary Figure 8). This performance benefit of conditional inference is due to the combination of less variable beliefs (and responses) at the end of ‘episodes’ – i.e., successive draws from the same bag – and equally fast adaptation to reversals (Supplementary Figure 8). By contrast, other belief stabilization strategies trade less variable responses against slower adaptation to reversals. A ‘soft’ version of the conditional inference model (see Methods) that down-weights the evidence that falls below the reliability threshold rather than discarding it also predicts an inverted U-shaped relation with accuracy. However, it fails at predicting a decreased behavioral variability and explains human behavior less accurately than the conditional inference model (exceedance *p* < 0.001; Supplementary Figures 9 and 10).

**Figure 8.**
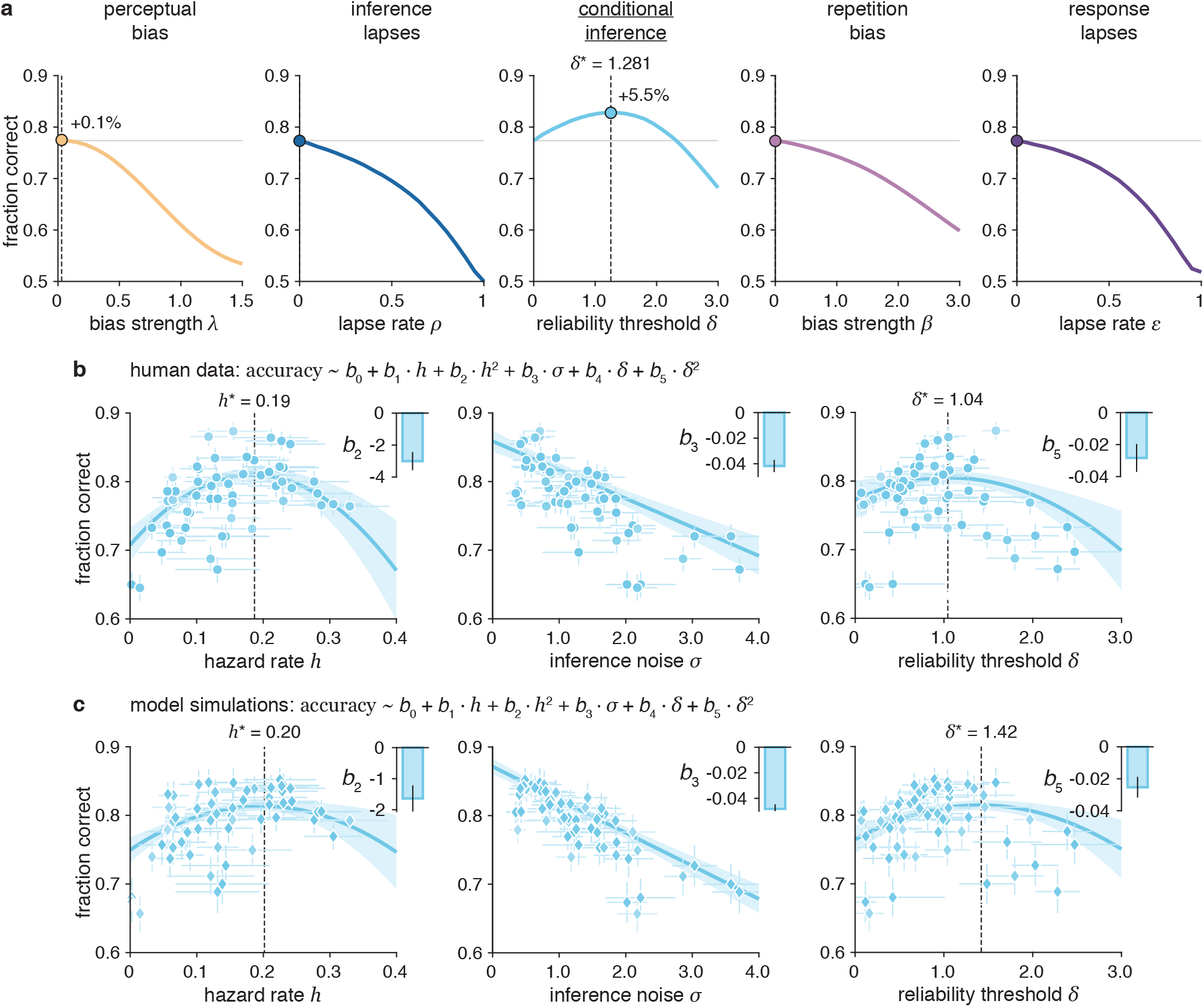
Increased decision accuracy through conditional inference. **(a)** Simulated effects of response stabilization strategies on decision accuracy. Fraction of overall correct responses when increasing each stabilization parameter drops for all models except for the conditional inference model, for which it increases the accuracy by +5.5% for *δ*\* = 1.28. Dots correspond to the best simulated accuracy. **(b)** Observed effects of best-fitting model parameters on human decision accuracy. For each best-fitting conditional inference model parameter, corresponding human accuracy (dots) and robust regression (solid blue lines, *r*-squared = 0.859, *p* < 0.001). Inverted U-shaped relation between reliability threshold and accuracy (negative quadratic coefficient *b*_5_, *p* = 0.002). Shaded area corresponds to the 95% confidence interval for predicted values. Parameters are presented as mean ± s.d. of posterior distributions and human accuracy as mean ± s.d. given the number of trials provided. Top right insets correspond to estimated regression coefficients presented as mean ± s.e.m. **(c)** Predicted effects of best-fitting model parameters on modeled decision accuracy. For each best-fitting conditional inference model parameter, corresponding conditional inference model predicted accuracy (diamonds) and robust regression (solid blue lines). Inverted U-shaped relation between reliability threshold and accuracy (negative quadratic coefficient *b*_5_, *p* < 0.001). Shaded areas correspond to the 95% confidence interval for predicted values. Parameters are presented as mean ± s.d. of posterior distributions and vertical error bars correspond to the accuracy s.d. of best-fitting model simulations. Top right insets correspond to estimated regression coefficients presented as mean ± s.e.m.

We measured this performance benefit of conditional inference in participants’ data by fitting a multiple regression model of accuracy as a function of each parameter: the hazard rate *h* (entered as a quadratic effect since it is expected to show an inverted U-shaped relation with accuracy, maximal accuracy expected around the true value of the hazard rate, see Methods), the inference noise *σ*_inf_ (entered as a linear effect, since it is expected to show a negative relation with accuracy), and the reliability threshold *δ* (entered as a quadratic effect since it is also expected to show an inverted U-shaped relation with accuracy; Figure 8a). This multiple regression model provides an accurate fit to participants’ data (Figure 8b; *r*-squared = 0.859, *d.f*. = 53, *p* < 0.001), and confirms the inverted U-shaped relation between reliability threshold and accuracy (quadratic coefficient *b*_5_ = 0.028 ± 0.009, mean ± sem, *t*_53_ = 3.2, *p* = 0.002, 95% CI = [−0.046 – 0.011]). The accuracy-optimizing value of the reliability threshold estimated from the best-fitting model (*δ*\* = 1.042±0.174, mean ± s.e.m.) is consistent with simulations (Figure 8a). We confirmed that the quadratic relation between reliability threshold observed in participants’ data is predicted by simulations of the best-fitting conditional inference model, by replacing participants’ accuracy with the accuracy of model simulations (Figure 8c; *b*_5_ = −0.025 ± 0.006, *t*_53_ = −3.9, *p* < 0.001, 95% CI = [−0.0383 – 0.0123]). Implementing conditional inference with the optimal reliability threshold *δ*\* is associated with comparable performance improvements in participants and simulations (participants: +3.2%; simulations: +5.5%). These findings confirm that conditional inference not only mitigates the variability of beliefs arising from imprecise inference, but with this improves the accuracy of these beliefs by trading the costs of inference against its expected benefits based on the reliability of incoming sensory information.

## Discussion

Human observers combine multiple pieces of sensory evidence to form or revise beliefs about the current uncertain state of their environment. These beliefs remain nonetheless imprecise, not only because of the expected uncertainty associated with each piece of evidence or the unexpected uncertainty associated with transitions between states [Soltani, 2019], but also due to the imprecision of the inference process itself [Drugowitsch, 2016; Wyart, 2016; Findling, 2021]. Variability in beliefs might therefore help adapt to new contingencies, but might also be mistaken for genuine changes in the environment and trigger unwarranted changes-of-mind. Here, we investigated whether and how human observers mitigate the trial-to-trial variability of their imprecise beliefs, by studying human behavior in a volatile decision-making task. Using computational modelling, we found that humans stabilize their imprecise beliefs using an efficient ‘conditional’ strategy – whereby they discard pieces of sensory evidence that are not deemed as sufficiently reliable. We show that this conditional belief updating strategy is associated with large individual differences linked to participants’ perceptual confidence, and improves the accuracy of resulting decisions.

It is well-known that human decisions show a suboptimal trial-to-trial variability under uncertainty [Wyart, 2016; Findling, 2021]. This behavioral variability can arise from internal ‘noise’ located at different stages of information processing. Classical cognitive models consider behavioral variability as a consequence of noise corrupting either sensory processing or the decision policy, in this later case by introducing a softmax at the action selection stage [Sutton, 1998; Griffiths, 2006; Glaze, 2015]. However, recent findings have shown that the large fraction of the observed behavioral variability arises from computation noise in inference [Drugowitsch, 2016; Findling, 2021; Findling, 2019]. Both policy noise and inference noise generate behavioral variability, but these two sources of noise differ regarding the trial-to-trial variability of underlying beliefs. Indeed, policy noise does not generate any trial-to-trial variability in beliefs but only variability in behavior. By contrast, inference noise generates trial-to-trial variability in both beliefs and behavior. Recent neurophysiological findings support the idea that the large moment-to-moment variability of neural activity in cortical areas which reflect evidence accumulation corresponds to genuine variability in beliefs [Peixoto, 2021].

Large moment-to-moment fluctuations in beliefs are likely to be costly, both psychologically and neurally. At the psychological level, changes-of-mind require changes in behavior, which entail increased cognitive control and slower response latencies. At the neural level, previous studies have shown that changes-of-mind are associated with strong increases in neural activity in the prefrontal cortex in humans [Donoso, 2014] and rodents [Karlsson, 2012]. We thus reasoned that humans may rely on a form of ‘belief stabilization’ strategy to mitigate the impact of inference noise on belief variability and prevent unwarranted changes-of-mind. We found that human behavior is less variable than what is predicted from the amount of inference noise that matches behavioral accuracy. To characterize how humans reduce the variability of their behavior, we compared several stabilization strategies at different processing stages. Some of them mitigate the variability of behavior but not beliefs, whereas others stabilize the underlying beliefs [Luu, 2018; Lange, 2021]. We found that human observers use a ‘conditional’ strategy to stabilize their beliefs. This means that humans discard (ignore) a fraction of the available sensory evidence judged as too weak and unreliable. Interestingly, this strategy challenges the idea that more information is always better in presence of inference noise when performing a volatile decision-making task.

Previously described strategies typically reduce the variability of beliefs through some form of ‘confirmation bias’, either by shifting the perception of sensory evidence in the direction of prior beliefs as in the ‘perceptual bias’ model that we have considered among alternative stabilization strategies, or by underweighting (or even discarding) the sensory evidence that is inconsistent with prior beliefs. In particular, recent work [Glickman, 2022; Salvador, 2022] has identified such a ‘stimulus-consistency’ bias mechanism during sequential cue combination. This bias, which resembles a similar bias described during reinforcement learning [Lefebvre, 2017], was observed in stable environments. In these conditions where the latent cause does not change, it was shown to increase accuracy by reducing decision sensitivity to misleading sensory observations. By contrast, in volatile environments, the same bias typically decreases accuracy because inconsistent observations can signal a change in their latent cause, and should therefore not be underweighted [Lefebvre, 2022]. Future work should further investigate the effect of task parameters on the stabilization strategies deployed by humans to mitigate the negative effects of inference noise.

In our task, tested participants discard about a third of presented stimuli, a surprisingly large fraction for stimuli categorized in isolation with 80% overall accuracy in a forced-choice task. Importantly, the ‘lapse’ rate measured in the condition where participants categorize isolated stimuli is widely smaller than the ‘conditional discard’ rate measured in the main task (lapse rate: 9.3%; conditional discard rate: 28.9%). This means that the conditional inference strategy observed in our task does not reflect lapses in attention. Consistently with the idea that apparent lapses in behavior do not only reflect inattention, recent work in mice has shown that lapses measured during perceptual decisions can reflect exploration [Pisupati, 2021] or alternations between discrete behavioral strategies [Ashwood, 2022]. In our task, we propose that the discarding of unreliable stimuli is controlled by a general metacognitive process linked to perceptual confidence. Indeed, the reliability threshold used for discarding stimuli correlates with the confidence threshold measured in the same participants during an independent task.

In addition, tethering inference to the reliability of incoming sensory evidence shows positive effects as it reduces the cognitive costs of imprecise inference, in terms of unwarranted changes-of-mind as already mentioned, but also in terms of mental effort by waiving inference on a substantial fraction of trials. The fact that mental effort can be aversive enough for people to accept non-negligible levels of physical pain [Vogel, 2020] can thus partially explain why humans rely on an efficient metacognitive strategy to reduce mental effort. We further show that conditional inference simultaneously increases the accuracy of decisions while decreasing mental effort, making it a win-win strategy in our standard reversal learning task.

Like most decision parameters [Moutoussis, 2021], conditional inference shows large individual differences across tested participants. We observed that the reliability threshold used to discard unreliable stimuli is associated with a specific behavioral gradient across participants. The fact that the position of individual participants on this gradient remains relatively stable across both experimental sessions indicates a genuine trait-like parameter. One important question concerns the relation between this trait-like parameter and other cognitive traits. We have shown that the reliability threshold used to discard stimuli correlates with the confidence threshold adopted by the same participants in a different task – another trait-like parameter which shows high test-retest reliability across experimental sessions. This means that the reliability threshold could be related to transdiagnostic psychiatric symptom dimensions [Gillan, 2016] known to correlate with confidence threshold in various tasks [Rouault, 2018].

In our task, we introduced expected uncertainty by degrading stimuli using sensory noise, but this is not the only way by which expected uncertainty can be introduced in decision tasks [Lange, 2021]. Indeed, existing tasks often introduce uncertainty by reducing the information conveyed by a single stimulus about its generative category – a form of uncertainty referred to as category ambiguity. The reliability-contingent control of inference observed in our task has not been identified in previous tasks where, unlike the present task, the sensory reliability of presented stimuli is large and it is category ambiguity that is low [Drugowitsch, 2016]. Future work could investigate more explicitly the different strategies deployed for mitigating the impact of imprecise beliefs in conditions of high sensory noise vs. high category ambiguity. Another important question concerns the neural processes that may underlie this conditional inference strategy. Our model is purely conceptual and suggests a form of hierarchical inference, in line with other proposed belief stabilization strategies [Luu, 2018; Lange, 2021]. In our model, this means that beliefs are updated downstream (and separately from) the categorization and evaluation of individual stimuli. Future work will be needed to investigate the possible neural mechanism underlying conditional inference, which may be a form of ‘gating’ – an all-or-none, non-linear filtering mechanism – that requires sufficient sensory response to trigger changes in the downstream decision (belief updating) circuit.

Taken together, our results show that humans condition the imprecise update of their beliefs to the reliability of incoming sensory information – an efficient metacognitive control process which not only prevents unwarranted changes-of-mind, but also reduces mental effort by waiving belief updating. Avoiding the mental effort spent on cognitive computations that are less likely to pay off (in this case, based on unreliable evidence) proves a surprisingly effective strategy that simultaneously increases the objective accuracy of resulting decisions – thereby bypassing the general cognitive trade-offs between the benefits and costs of cognitive operations described in the literature [Shenhav, 2013].

## Methods

### Participants

Sixty healthy adult participants (30 females, age: 25.8±3.9 years, all right-handed, Experiment 1: *N* = 30, 15 females, age: 24.7 ± 3.9 years; Experiment 2: *N* = 30, 15 females, age: 27.16 ± 3.7 years) took part in the study. One participant was excluded from analyses due to poor performance (overall decision accuracy more than 3.5 s.d. below the group-level mean). Participants were screened for the absence of any history of neurological and psychiatric disease or any current psychiatric medication. All participants provided written informed consent, and received 45 euros as compensation for their participation after completing the second experimental session. The study followed guidelines from the Declaration of Helsinki and specific procedures which received ethical approval from relevant authorities (Comité de Protection des Personnes Ile-de-France VI, ID RCB: 2007-A01125-48, 2017-A01778-45).

### Task and procedure

Both experiments were based on the same reversal learning task. In repeated trials, participants were presented with visual cues described as ‘marbles’ which could be drawn either from a bag containing light marbles or a bag containing dark marbles. Participants had to infer the bag the marble was drawn from (light or dark). Participants were instructed that marbles were not drawn randomly and independently across trials, but from the same bag for a certain number of successive trials (i.e., an episode) before being drawn from the other bag (i.e., a generative category reversal). Participants were herewith instructed about the presence of reversal but did not know the number of trials (or episode length) before a reversal. The length of these episodes was drawn from a bounded exponential distribution, resulting in an approximately flat hazard rate – a procedure equivalent to a Markov process with fixed transition probability. Each marble corresponded to a two-tone disc with light and dark shades of gray, in different proportions and spatially scrambled over the surface of the disc. The disc was generated using Gaussian noise filtered through a two-dimensional Gaussian smoothing filter and eventually binarized accordingly to the expected proportions between the light and dark shades of gray. All stimuli were presented on a uniform medium gray background. In experiment 1, the relative proportions of light and dark shades of gray were adjusted using a titration procedure to reach 80% of marbles perceived as light for the light bag, and 80% of marbles perceived as dark for the dark bag. In experiment 2, three levels of difficulty were titrated for each bag to achieve correct stimulus identification of 70%, 80% and 90%, respectively. In this second experiment, participants were not explicitly instructed of the 3 difficulty levels.

The titration procedure preceded each reversal learning block, and corresponded to two adaptive psychophysical staircases [Kaernbach, 1991] adjusting the relative proportions of light and dark shades of gray in presented marbles. We ran two separate staircases because no symmetry was assumed: participants could have a biased perception towards the light or the dark category. The trials of the two staircases were randomly interleaved during the procedure. In experiment 1, each staircase converged toward a light/dark proportion, one proportion was used to generate the marbles populating the light and the other for marbles populating the dark bag, each corresponding to 80% accuracy (light bag: 52.3 ± 1.1%, dark bag: 47.6 ± 1.0%). In experiment 2, an additional logistic regression was performed to extrapolate the light/dark proportions corresponding to 70% and 90% accuracy levels from the light/dark proportions corresponding to 80% accuracy (light bag: 51.6 ± 1.4% for 70% accuracy, 52.6 ± 1.5% for 80% accuracy, 54.1 ± 1.7% for 90% accuracy; dark bag: 48.40 ± 1.3% for 70% accuracy, 47.4 ± 1.4% for 80% accuracy, 45.9 ± 1.5% for 90% accuracy). The titration blocks were presented to participants as a distinct game (‘game #1’): participants were instructed to sort individual marbles, and informed that marbles were presented in random order (unlike the main task). Auditory feedback was provided at the end of each trial in titration blocks.

The reversal learning blocks were presented as ‘game #2’ with instructions to report whether the computer is drawing marbles from the light or the dark bag after the presentation of each marble. Unlike the titration blocks, no feedback was provided during the reversal learning blocks. The experiment consisted of 16 blocks (8 blocks from the task condition described above, and 8 blocks of another task condition not relevant for the current study), divided in two experimental sessions of 8 blocks, each session lasting approximately 90 min and taking place on different days. Each reversal learning block consisted of 80 trials. Experiment 1 consisted of two types of blocks: ‘low volatility’ blocks containing 5 reversals (i.e., hazard rate *h* = 1/16) and ‘high volatility’ blocks containing 10 reversals (i.e., hazard rate *h* = 1/8). Experiment 2 consisted of ‘high volatility’ blocks only. The first generative category of each block was counter-balanced pseudo-randomly across blocks and participants.

Participants were instructed that each marble consisted of both light and dark shades of grey spatially scrambled. They were also instructed that all marbles contained in the light bag were predominantly light, whereas all marbles contained in the dark bag were predominantly dark (Supplementary Figure 12). Before the experiment, they performed a short training for both ‘games’ (titration and reversal learning blocks) to get familiarized with the stimuli and the difficulty of identifying this predominant dark/light color due to the spatial scrambling and the short stimulus presentation time (83 ms).

The experiment was coded in MATLAB and run using Psychtoolbox-3 [Kleiner, 2007]. Participants performed the experiment in a soundproof booth with their head positioned on a chin rest at 75 cm from a 24-inch LCD screen operating at 60 Hz with a resolution of 1920 × 1080 pixels.

### Bayes-optimal inference

For each trial *t*, the decision-maker tries to find the associated hidden state corresponding to one of the two alternative categories *c_t_* = +1/ – 1 (light or dark bag). At each trial, the belief about the current value of the hidden state is updated by combining the prior belief with the new incoming evidence according to Bayes rule. The normalized model of decisions between two alternatives is formalized in [Glaze, 2015] who define the belief at trial *t*, *L_t_*, as the logarithm of the posterior odds of the alternative categories accumulated until this trial. The sign of the log-odds belief indicates which category is more likely, whereas the magnitude of the log-odds belief indicates the strength of the belief in favor of the more likely hidden state. The update rule combining prior belief *L*_*t*–1_ and new incoming evidence *LLR_t_* is defined as follows:

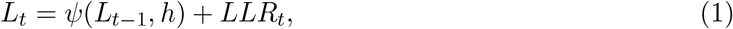

with *ψ* the time-varying prior expectation defined by

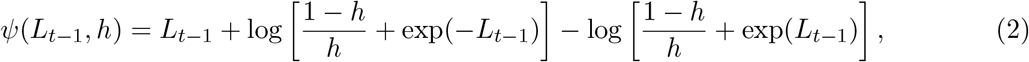

and *h* the hazard rate, the expected probability at each trial t that the category will switch, which corresponds to a non-linear integration leak. We also considered a linear approximation of the prior expectation function [Glaze, 2015] (Eq. 2) turning the model into a leaky evidence accumulator (Supplementary Figure 2):

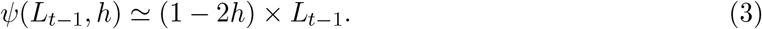

#### Suboptimal Bayesian inference

Each update of the log-odds belief with new stochastic incoming evidence *LLR_t_* is corrupted by internal gaussian noise – inference noise – with standard deviation *σ*_inf_ turning the deterministic belief update equation (Eq. 1) into a stochastic draw [Drugowitsch, 2016; Weiss, 2021]:

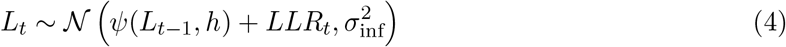

#### Model-based analysis of behavior and stabilization strategies

We modeled human behavior in the experimental condition using a hierarchical suboptimal Bayesian model with two free parameters *h* and *σ*_inf_ and with an additional model-specific stabilization parameter. We consider the sensory response to the stimulus presented at trial t as a normally distributed random variable generated by 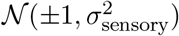, depending on the true underlying category of the presented stimulus (+1 for light, −1 for dark). The standard deviation of sensory noise *σ*_sensory_ is set to correspond to 80% of correct categorization – achieved through the online titration procedure.

In the first (‘categorized’) version of the model, this sensory response is then categorized as light (*c_t_* = +1) or dark (*c_t_* = −1) and expressed as a log-odds ratio. By design of the task, the log-likelihood ratio is ideally 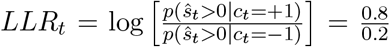 for evidences towards *c_t_* = +1 and respectively 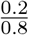 for evidences towards *c_t_* = −1. In the second (‘non-categorized’) version of the model, the log-odds *LLR_t_* is computed using the continuous normally distributed stimulus sensory response without any explicit categorization (Supplementary Figure 9). The log-odds ratio between two symmetrical gaussian distributions 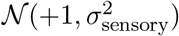 and 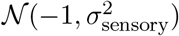 or vice-versa, simplifies to a scaled version of the sensory response by a factor 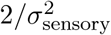 thus normally distributed with mean equal to 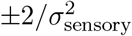 and s.d. equal to 2/*σ*_sensory_. The combination of the evidence provided by the stimulus (*LLR_t_*) and the prior belief (*LLR*_*t*–1_, also expressed as log-odds ratio and updated according to hazard rate *h*) is then corrupted by Gaussian inference noise of standard deviation *σ*_inf_ following Eq. 4. Inference noise captures both internal variability in the perceived log-odds ratio *LLR_t_* and variability in the update of the belief *L_t_*.

The response selection policy is based on the sign of the decision variable: the newly formed belief *L_t_*. We considered a normative, ‘greedy’ selection process and a noisy selection process (Supplementary Figure 1) modeled by sampling responses from the sign of a normally distributed decision variable with mean *L_t_* and variance 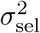.

We introduced stabilizing parameters to the suboptimal Bayesian inference process described above, operating at different stages of information processing.

1. The perceptual bias model, controlled by a prior bias strength λ, shifts the sensory representation of the stimulus already expressed as log-odds 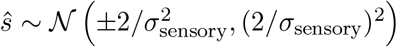 towards the prior belief *L*_*t*–1_:

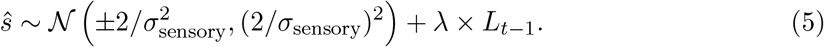 In this case, the log-likelihood ratio differs from ideal as the integration range is shifted by λ × *L*_*t*–1_.
2. The inference lapse model, controlled by a fraction of lapses *ρ*_lapse_, ignores a fraction of the stimuli whose evidence is not used to update the current belief:

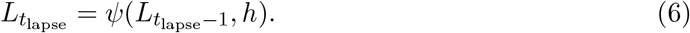
3. The conditional inference model, controlled by a reliability threshold *δ*, uses only the stimuli whose sensory responses exceed a fixed threshold, |*ŝ*| > *δ*, to update the current belief:

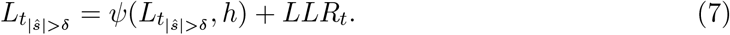 In this case, the log-likelihood ratio *LLR_t_* accounts for the presence of the reliability threshold by integrating conditional likelihoods above the reliability threshold (instead of zero for the standard model). If the sensory response does not exceed the reliability threshold, the stimulus is ignored, leading to:

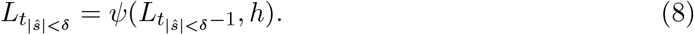 A discard rate – corresponding to the fraction of unreliable evidence not used for belief update – can be derived by integrating the truncated cumulative distribution function 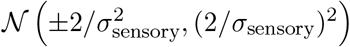 on the remaining range [−*δ*; +*δ*].
4. The confirmation bias model, controlled by a bias strength *β*, operates at the selection stage by systematically shifting the decision criterion in favor of the previous response:

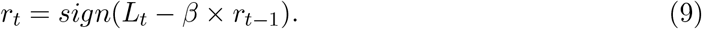
5. The response lapse model, controlled by a lapse rate *ϵ*, makes a fraction of ‘blind’ repetitions of the previous response irrespective of the current belief:

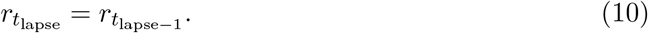 We considered a ‘soft’ version of the conditional inference model (Supplementary Figures 9 and 10), controlled by the same reliability threshold *δ*, not discarding but down-weighting the stimuli whose sensory responses do not exceed the threshold. In this model, inference is performed at every trial but with two distinctly weighted log-likelihood ratio *LLR_t_*, the first integrating likelihoods below and the second integrating likelihoods above the reliability threshold.

### Fitting procedure

Due to the presence of internal noise during sensory processing and hidden-state inference, which propagates across trials, no solution for model likelihoods can be derived analytically. We therefore used simulation-based methods to obtain noisy likelihoods for tested sets of parameter values, which were fed to specific fitting algorithms capable of handling such noisy likelihood functions.

Instead of fitting the candidate models to all choices using simulation-based methods, we focused on the two behavioral signatures characteristic of reversal learning behavior: the response reversal curve and the response switch curve. This focused fitting procedure aimed at testing which of the tested candidate models can best explain specifically our behavioral effects of interest. Note that fits to all choices yield the same results (Supplementary Figure 11). We fitted the parameter values (*h*, *σ*_inf_ and the optional stabilizing parameter) which best explained these two psychometric curves, both evaluated on 4 trials preceding and 4 trials following a reversal, for a total of 8 trials per reversal for each metric. We conducted model recovery analyses, detailed below, to ensure the reliability of the simulation-based fitting method outlined above.

In practice, we simulated model responses (*N* = 1000) to the sequences of trials presented to participants, we then derived the corresponding mean and standard deviation of reversal and switch curves (modeled as normally distributed variables at each trial) and estimated the log-likelihood of observed (participant) reversal and switch curves given model simulations. We combine the loglikelihood estimate with prior distributions on parameter values to get a log-posterior estimate, whose prior distributions over parameters were defined as truncated beta and gamma distributions (*h*: beta distribution with shape parameters *α* = 2 and *β* = 18, range [0, 0.5]; *σ*_inf_: gamma distribution with shape parameter *k* = 4 and scale parameter *θ* = 0.25, range [0, 5]; λ and *δ*: gamma distribution with shape parameter *k* = 1 and scale parameter θ = 0.75, range [0, 5]; *ρ*_lapse_ and *ϵ*: beta distribution with shape parameters *α* = 1 and *β* = 9, range [0, 1]; *β*: gamma distribution with shape parameter *k* = 1 and scale parameter *θ* = 0.25, range [0, 5]). In the first experiment, we verified for the winning model that using looser prior distributions did not change the best-fitting parameter values (Spearman correlations: *h*: *ρ* = 0.779, *p* < 0.001, 95% CI = [0.6500.784]; *σ*_inf_: *ρ* = 0.854, *p* < 0.001, 95% CI = [0.7630.862]; *δ*: *ρ* = 0.834, *p* < 0.001, 95% CI = [0.6700.843]; differences: *h*: *t*_28_ = 0.508, *p* = 0.615, Cohen’s *d* = 0.094, 95% CI = [−0.022+0.013], *BF*_01_ = 4.497; *σ*_inf_: *t*_28_ = 0.863, *p* = 0.395, Cohen’s *d* = 0.160, 95% CI = [−0.349 + 0.142], *BF*_01_ = 3.603; *δ*: *t*_28_ = 1.152, *p* = 0.259, Cohen’s *d* = 0.214, 95% CI = [−0.326 + 0.091], *BF*_01_ = 2.778).

The unnormalized log-posterior is given as argument for the Variational Bayesian Monte Carlo (VBMC) algorithm [Acerbi, 2020] (version 1.0; https://github.com/lacerbi/vbmc) returning a variational approximation of the full posterior and a lower bound on the log-marginal likelihood. The VBMC algorithm supports stochastic estimates of the log-posterior and we provide estimates of its standard deviation by bootstrapping the standard deviation of the estimated log-likelihood. We finally take the posterior mean to obtain best-fitting parameter values.

For the second experiment, we computed 3 distinct reversal curves and 3 distinct switch curves depending on the strength of the stimulus presented on the first trial following a reversal, for both model simulations and human data. We summed the log-likelihood across these 6 metrics.

We used the lower bound of the marginal likelihood as model evidence for Bayesian Model Selection (BMS) analysis. BMS was conducted using a random-effects approaches assuming that different participants may rely on different models, and consists in estimating the distribution over models that participants draw from. We used the Dirichlet parameterization of the random-effects approach implemented in SPM12 [Stephan, 2009; Rigoux, 2014] with default Dirichlet prior set to one – corresponding to a uniform distribution (http://www.fil.ion.ucl.ac.uk/spm).

Our approach focusses on behavior around reversals through the two psychometric curves described above. Alternatively, we considered fitting human data using per-trial choice log-likelihoods using a particle filter and the same VBMC algorithm. In this case, each trial – and not only trials around reversals – equally contributes to the goodness of fit, rather than focusing on our behavioral effects of interest around reversals. Yet, the more responses considered, the more likely a response can reflect a lapse and be a blind repetition of the previous response. To prevent the fitting procedure to try to fit those responses with the same importance as the other responses – especially the ones around reversal –, ergo to prevent those lapse responses to corrupt the goodness of fit, we included a lapse rate parameter *ϵ* as additional parameter to each first four stabilizing models (the fifth previously described candidate model being fully formalized by the lapse rate parameter *ϵ*). The prior distributions on parameter values remain the same as previously reported, except for the additional parameter *ϵ* meant to be kept low (beta distribution with shape parameters *α* = 1 and *β* = 39, range [0, 1]). The analysis of model fits to all choices using the same BMS yielded the same results as the ones obtained from model fits to behavioral effects of interest using our simulation-based approach (Supplementary Figure 11).

### Model recovery procedure

Model simulations were done for each participants’ true stimulus sequences using parameters corresponding to the mean of their respective prior distributions. We used prior distribution in order to have an *ex-ante* assessment that the constructed models are well-posed. Each simulation was fitted by a given model and we used the Dirichlet parameterization as described above to estimate the distribution over recovered models. We then sampled 10^6^ times from the estimated Dirichlet distributions and computed column-wise (for each recovered model) the exceedance probability of each simulated model.

### Confidence fitting procedure

We modeled human behavior in titration trials using three free parameters, the sensory noise s.d. *σ*_sensory_, the confidence noise s.d. *σ*_confidence_ and the confidence threshold *δ*_confidence_. Each stimulus sensory response at trial t is modeled as a normally distributed random variable generated by 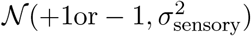 depending on the staircase it originates from. Based on this sensory response, the noisy confidence is reported with respect to a confidence threshold*δ*_confidence_. We fitted this model using a particle filter (*N* = 1000) to the factorized responses to titration trials (light and unconfident; light and confident; dark and unconfident; dark and confident) using VBMC to maximize the log-likelihood of factorized response probabilities.

### Principal Component Analysis

Principal component analysis (PCA) was performed on participants’ response reversal and response switch curves using MATLAB in-built function which centers the data and uses the singular value decomposition (SVD) algorithm. Based on the first three obtained PC scores, we estimated the explained variance in reversal and switch curves through bootstrapping (*N* = 10, 000).

### Multiple Regression Model

We predicted human and conditional inference model accuracies as function of each parameter using a multiple regression model. We chose to include the perceived hazard rate *h* and the reliability threshold *δ* up to the second-order in the multiple regression, but the inference noise *σ_inf_* appears only up to the first-order as it only corrupts accumulation of evidence. Second-order dependencies, if any, allow to exhibit inverted U-shaped relations with accuracy – ergo a beneficial effect on accuracy. As we investigate the benefits of the conditional inference strategy, the reliability threshold *δ* was included up to the second-order, and showed indeed an inverted U-shaped relation with accuracy, no discarding of evidence corresponding to a classical suboptimal inference model and a large *δ* implying omission of almost all evidence. The multiple regression model was fitted using the MATLAB in-built function `fitnlm.m` with robust fitting (weighted least-squares).

### Statistical testing and reproducibility

Unless noted otherwise, statistical analyses of differences between scalar metrics in both experiments relied on two-tailed parametric tests (classical and Bayesian paired *t*-tests, repeated-measures ANOVA) between tested participants and model predictions in a paired manner using MATLAB in-built functions, `simple_mixed_anova.m` [Caplette, 2022] and JASP [JASP Team, 2021]. For repeated-measures ANOVA, reported *p*-values are Greenhouse-Geisser corrected. Given our sample sizes, these statistical tests were applied outside the small-sample regime. Data were not explicitly tested for normality. Reported statistics are not corrected for multiple comparisons. Given the absence of prior effect sizes, we chose a sample size (*N* = 30 for each experiment) which exceeded the average sample size used in human psychophysical studies with similar trial number per participant (*n* = 640 across two sessions).

## Data availability

The datasets generated during and analyzed during the current study is freely available online on figshare: https://figshare.com/projects/Efficient_stabilization_of_imprecise_statistical_inference_through_conditional_belief_updating/140170

## Code availability

The analysis code supporting the reported findings is freely available online on github: https://github.com/juliedrevet/CONDINF

## Acknowledgements

This work was supported by a starting grant from the European Research Council (ERC-StG-759341) awarded to Valentin Wyart, a US-France collaborative research grant from the National Institute of Mental Health (1R01MH115554-01) and the Agence Nationale de la Recherche (ANR-17-NEUC-0001-02) awarded to Jan Drugowitsch and Valentin Wyart, and a department-wide grant from the Agence Nationale de la Recherche (ANR-17-EURE-0017). We thank Benedetto De Martino and Marius Usher for their insightful comments and suggestions during peer review. The funders had no role in study design, data collection and analysis, decision to publish or preparation of the manuscript.

## Author contributions

**Julie Drevet**: Conceptualization, Methodology, Software, Validation, Formal Analysis, Investigation, Resources, Data Curation, Writing – Original Draft; Writing - Review & Editing; Visualization. **Jan Drugowitsch**: Conceptualization, Methodology, Writing - Review & Editing, Supervision, Project administration, Funding acquisition. **Valentin Wyart**: Conceptualization, Methodology, Software, Validation, Formal Analysis, Resources, Writing – Original Draft, Writing - Review & Editing, Visualization, Supervision, Project administration, Funding acquisition

