## Supplementary Information for "Efficient stabilization of imprecise statistical inference through conditional belief updating"

July 4, 2022

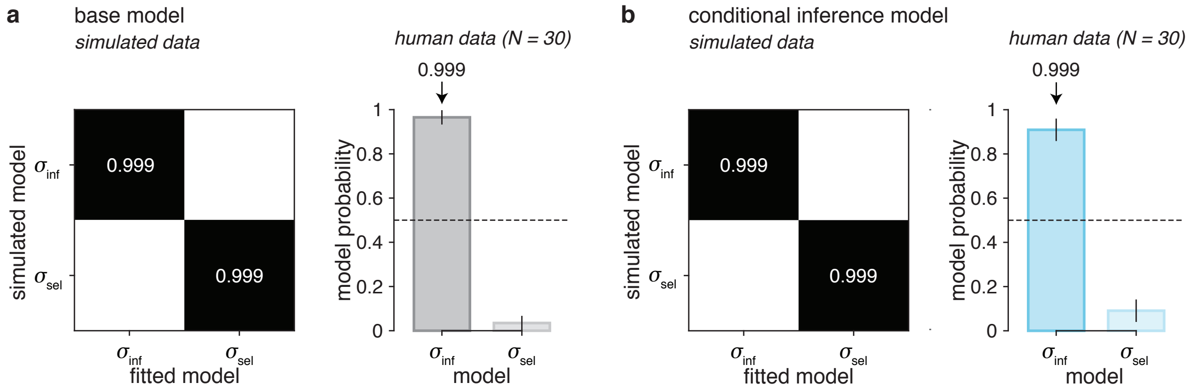


**Supplementary Figure 1 | Comparison between inference and selection noise. (a)** Bayesian inference model corrupted by internal noise (either inference noise or selection noise). Left: *ex-ante* confusion matrix from model recovery on simulated data, indicating that for each recovering noise source, the model used to simulate the data was the same with exceedance $p_{\mathrm{exc}} > 0.999$. Right: estimated model probabilities for inference and selection noise when fitted to human data, Bayesian model selection shows a clear preference for inference noise over selection noise with exceedance probability $p_{\mathrm{exc}} > 0.999$. Model probabilities are presented as mean and s.d. of the estimated Dirichlet distribution. **(b)** Conditional inference model. Left: *ex-ante* confusion matrix from model recovery on simulated data, indicating that for each recovering noise source, the model used to simulate the data was the same with exceedance $p_{\mathrm{exc}} > 0.999$. Right: estimated model probabilities for inference and selection noise when fitted to human data, Bayesian model selection shows a clear preference for inference noise over selection noise with exceedance probability $p_{\mathrm{exc}} > 0.999$. Model probabilities are presented as mean and s.d. of the estimated Dirichlet distribution.

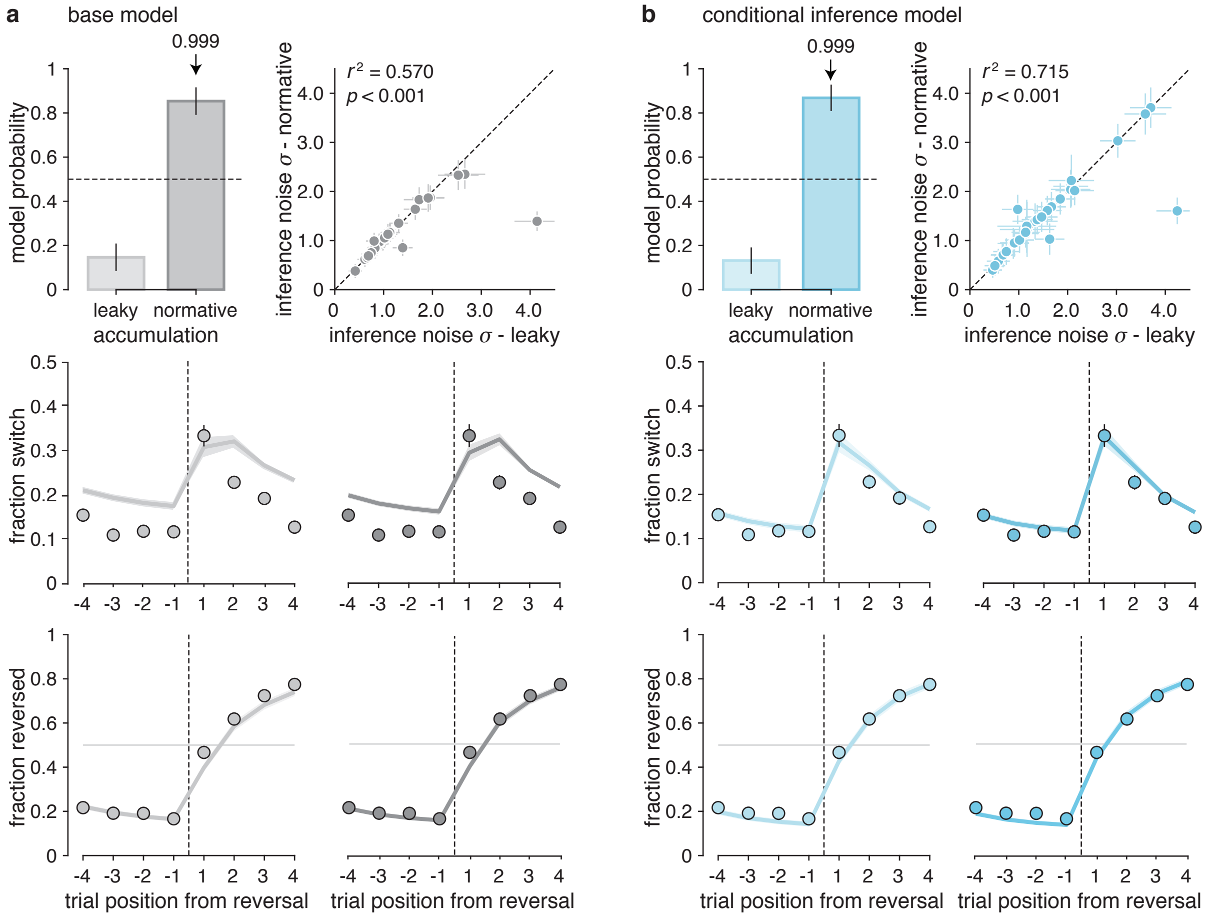


**Supplementary Figure 2 | Comparison between leaky and normative evidence accumulation** ($N = 30$). **(a)** Inference noise added to a leaky accumulation model or to the normative accumulation model. Top left: estimated model probabilities for leaky and normative accumulation when fitted to human data, Bayesian model selection shows a clear preference for normative over leaky accumulation with exceedance probability $p_{\text{exc}} > 0.999$. Model probabilities are presented as mean and s.d. of the estimated Dirichlet distribution. Top right: positive correlation between best-fitting inference noise parameter added to the leaky accumulation model and best-fitting inference noise parameter added to the normative accumulation model. Error bars correspond to posterior s.d. Bottom: Response reversal curves (top row) and response switch curves (bottom row) predicted by the best-fitting leaky accumulation model (light grey line) and best-fitting normative accumulation model (dark grey line). Both leaky and normative accumulation models capture correctly the accuracy of behavior surrounding reversals but overestimates the variability of the same behavior, especially the leaky accumulation model. The reversal is represented by the thin dotted line. Dots indicate human data (group-level average). Error bars and shaded areas correspond to s.e.m. **(b)** Same panels as in (a) but with additional conditional inference stabilization strategy (either leaky accumulation or normative accumulation). Similar amount of inference noise is found in both accumulation models.

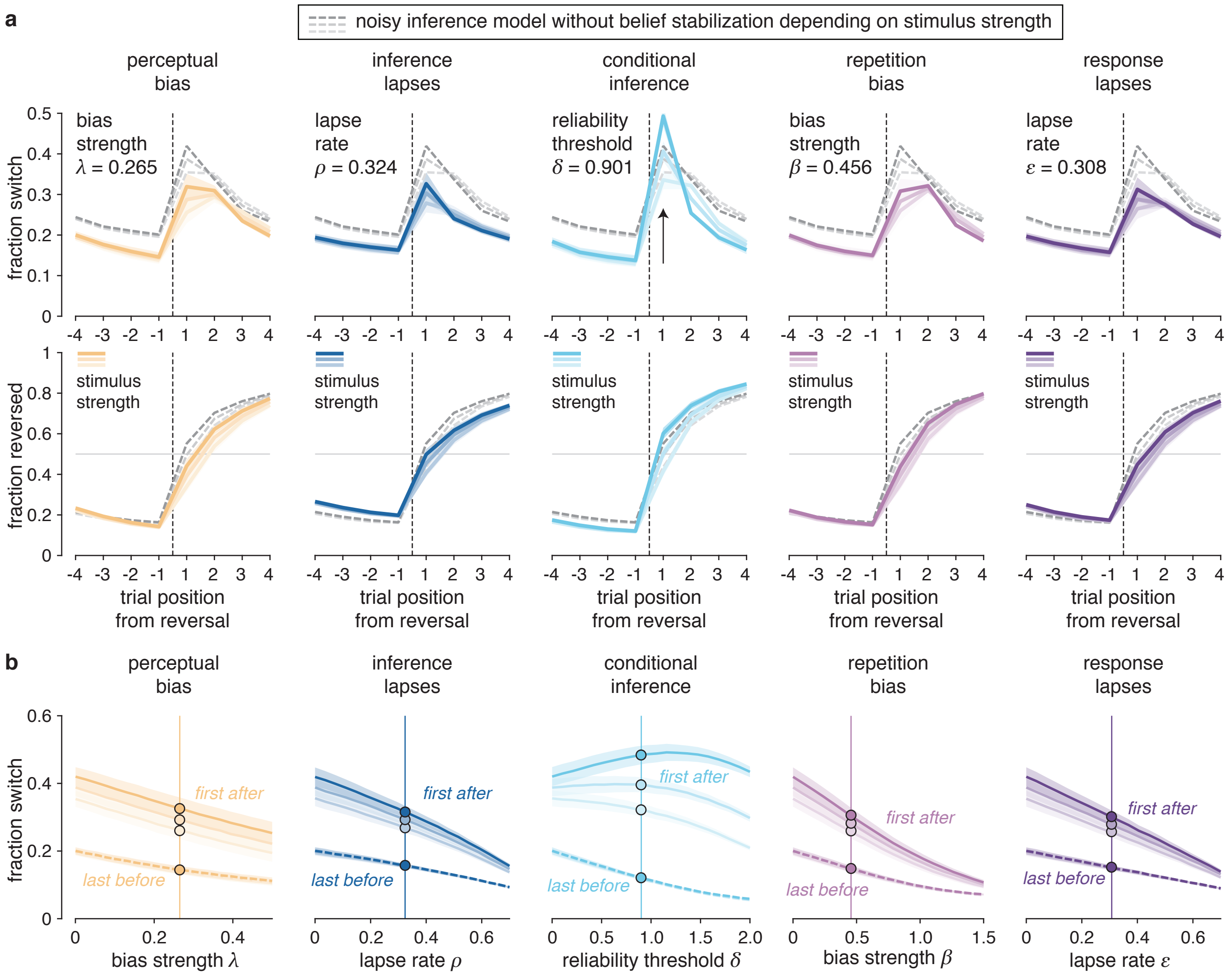


**Supplementary Figure 3 | Predicted effects of response stabilization strategies. (a)** Simulated effects of the five candidate strategies on response switch curves (top row) and response reversal curves (bottom row). The parameters controlling each response stabilization strategy ($\lambda$,$\delta$, $\rho$, $\beta$, $\epsilon$) is set to match participants' overall switch rate. The other parameters ($h$,$\sigma_{\text{sen}}$, $\sigma_{\text{inf}}$, $\sigma_{\text{sel}}$) are fixed to their best-fitting values for the noisy inference model without response stabilization. Solid colored lines correspond to the reversal behavior of each candidate model depending on the stimulus strength in the trial following a reversal. Dotted gray lines correspond to the reversal behavior of the noisy inference model without response stabilization depending on the stimulus strength in the trial following a reversal (same for all panels). Stronger color corresponds to stronger stimuli. The conditional inference strategy stands out from other strategies on the first trial after reversal (arrow). Shaded areas around curves correspond to s.e.m. (b) Simulated effects of the five candidate strategies on response switches just before and after a reversal. Fraction of response switches on the last trial before each reversal (dotted lines) and the first trial after each reversal (solid lines) for each belief stabilization strategy. Stronger colors correspond to a stronger (more informative) stimulus on the first trial following a reversal. All candidate strategies reduce simultaneously response switches before and after reversals, except for conditional inference which reduces response switches only before reversals (i.e., when they are not warranted). Dots correspond to stabilization parameters set as in (a). Shaded areas correspond to s.e.m.

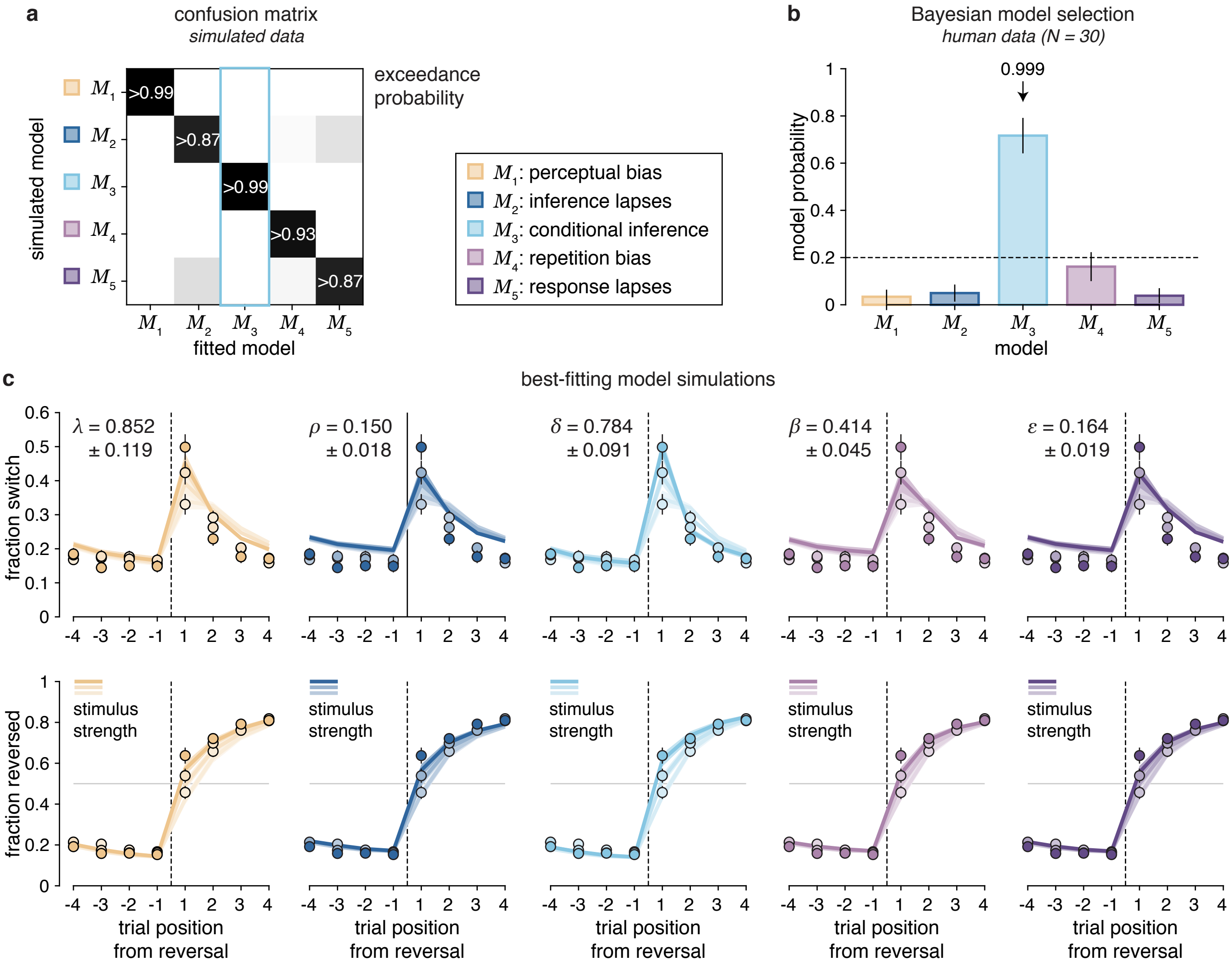


**Supplementary Figure 4 | Belief stabilization through conditional inference.** **(a)** Confusion matrix for *ex-ante* model recovery. For each recovering model, the model used to simulate the data was the same with exceedance $p_{\text{exc}} > 0.87$ and especially for the conditional inference model the exceedance reached is $p_{\text{exc}} > 0.999$. **(b)** Bayesian model selection of response stabilization strategy. Estimated model probabilities for the five candidate models. The conditional inference model outperforms other models with exceedance $p_{\text{exc}} > 0.999$. Model probabilities are presented as mean and s.d. of the estimated Dirichlet distribution. The dashed line corresponds to the uniform distribution. **(c)** Simulations of response stabilization strategies fitted to the human data. Simulations of three switch curves (top row) and three reversal curves (bottom row) are based on best fitting parameters for each stabilizing model. Dots indicate human data (group-level average), whereas lines indicate model predictions. The mean best-fitting parameter of each model (mean ± s.e.m.) is indicated in the top-left corner of each subpanel. Stronger colors correspond to a stronger (more informative) stimulus on the first trial following a reversal. Only the conditional inference model qualitatively reproduces humans stabilized behavior around reversal and the large transient increase of switches on the trial just after reversal. Shaded areas and error bars correspond to s.e.m.

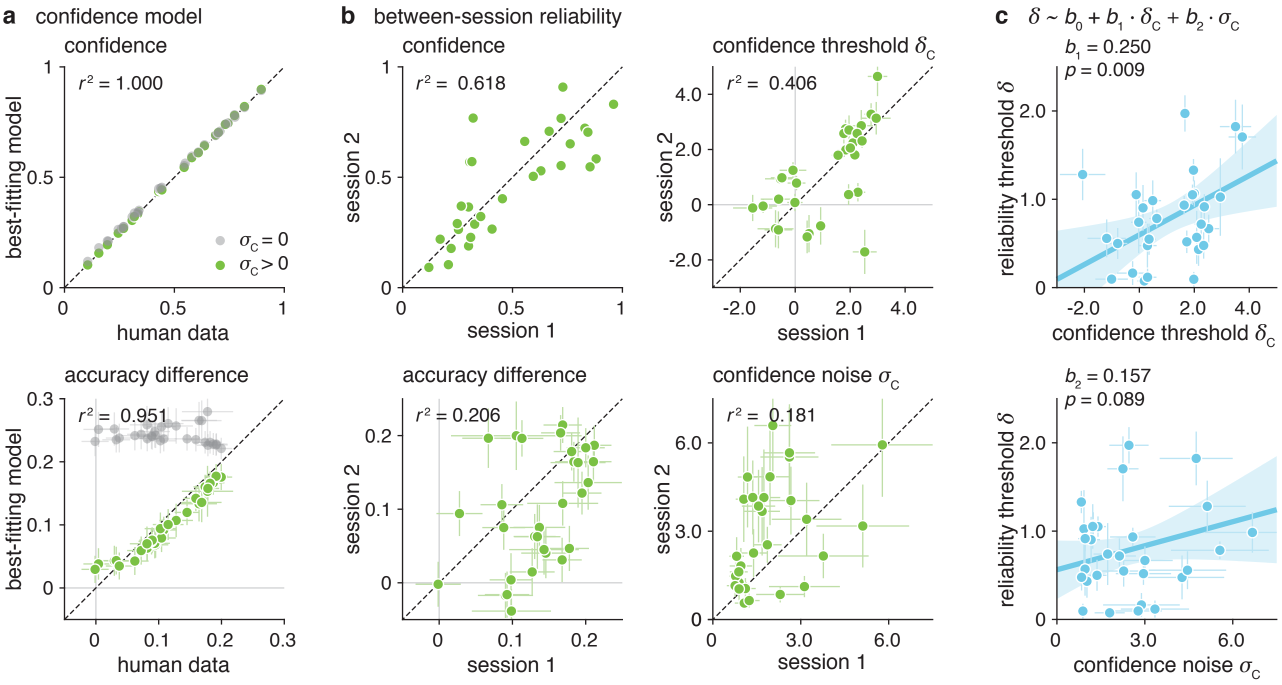


**Supplementary Figure 5 | Decision confidence model** ($N = 30$). **(a)** Reliability of confidence model predictions. Correlation between observed data and predicted confidence (top) and predicted accuracy difference between confident and unconfident trials (bottom) of a confidence model with (green) or without (gray) confidence noise $\sigma_c$. All data are presented as mean values and error bars correspond to s.d. of the accuracy difference of best-fitting model simulations and estimated s.d. of human accuracy difference given the number of trials provided. **(b)** Between-session reliability in confidence reports and parameters. Reported confidence and best-fitting parameters found for the first experimental session correlate positively with reported confidence and best-fitting parameters found for the second session. Data are presented as mean values. Error bars on best-fitting parameters correspond to s.d. of posterior distribution and error bars on accuracy difference correspond to estimated s.d. of human accuracy difference given the number of trials provided. **(c)** Reliability of positive relation between reliability threshold and confidence threshold. Linear Regression between reliability threshold and confidence threshold accounting for internal variability in confidence reports. Parameters are presented as mean ± s.d. of posterior distributions of each fit. Solid blue line correspond to the best-fitting regression line (in the least-squares sense) and shaded area corresponds to the 95% confidence interval for the regression line.

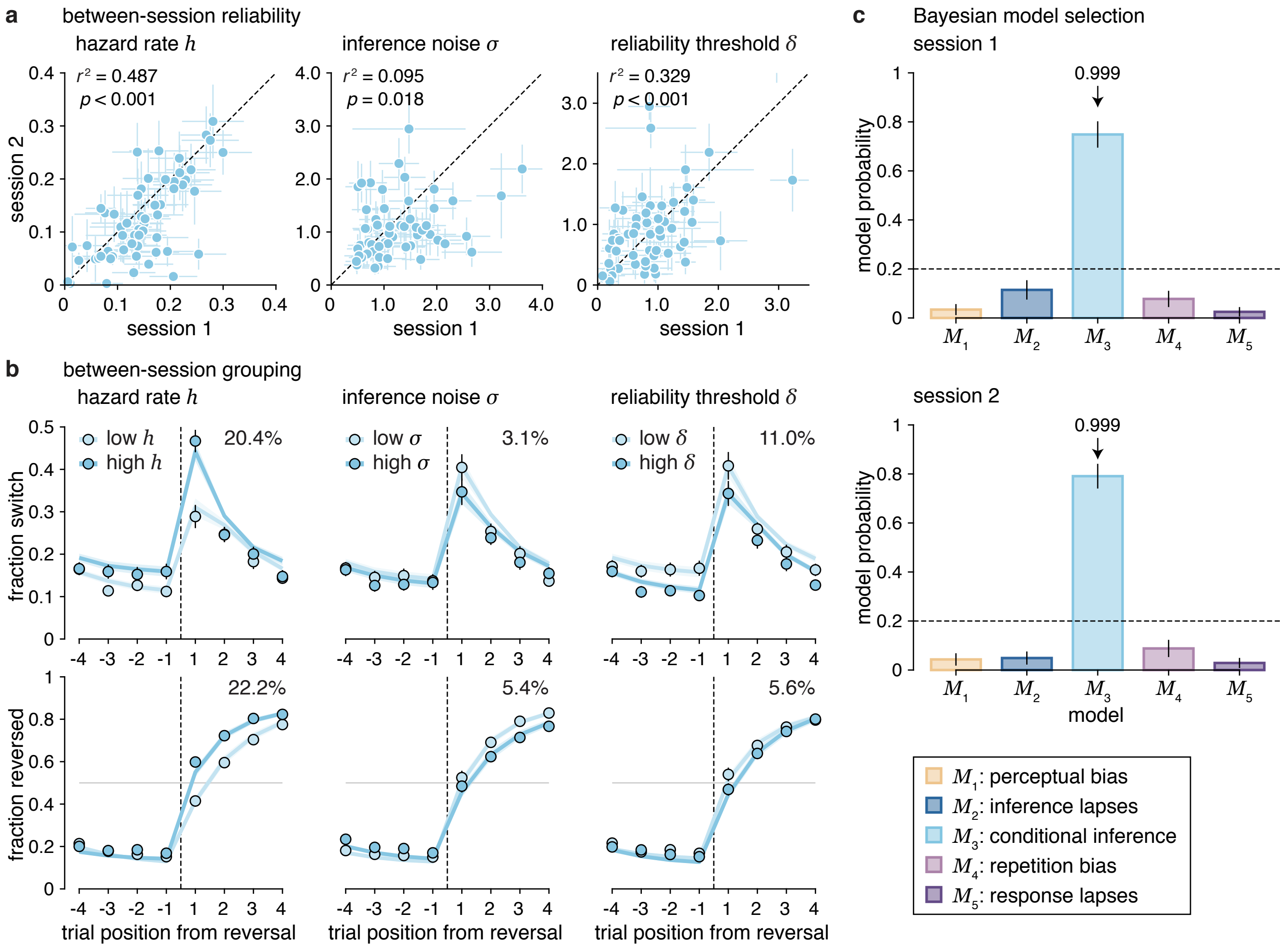


**Supplementary Figure 6 | Between-session reliability in conditional inference model parameters ($N = 60$).** **(a)** Between-session reliability in model parameter values. Best-fitting parameters found for the first experimental session correlate positively with best-fitting parameters found for the second session. Parameters are presented as mean ± s.d. of posterior distributions of each fit. Dotted lines correspond to the identity line. **(b)** Between-session (cross-validated) effect on response switch and reversal curves. Median-split on each best-fitting parameter for conditional inference model simulations (lines) and observed data (dots) and corresponding variance explained by each parameter. Error bars and shaded areas correspond to s.e.m. **(c)** Session- wise Bayesian model selection. The conditional inference model outperforms other models for both sessions with exceedance $p_{\mathrm{exc}} > 0.999$. Model probabilities are presented as mean and s.d. of the estimated Dirichlet distribution.

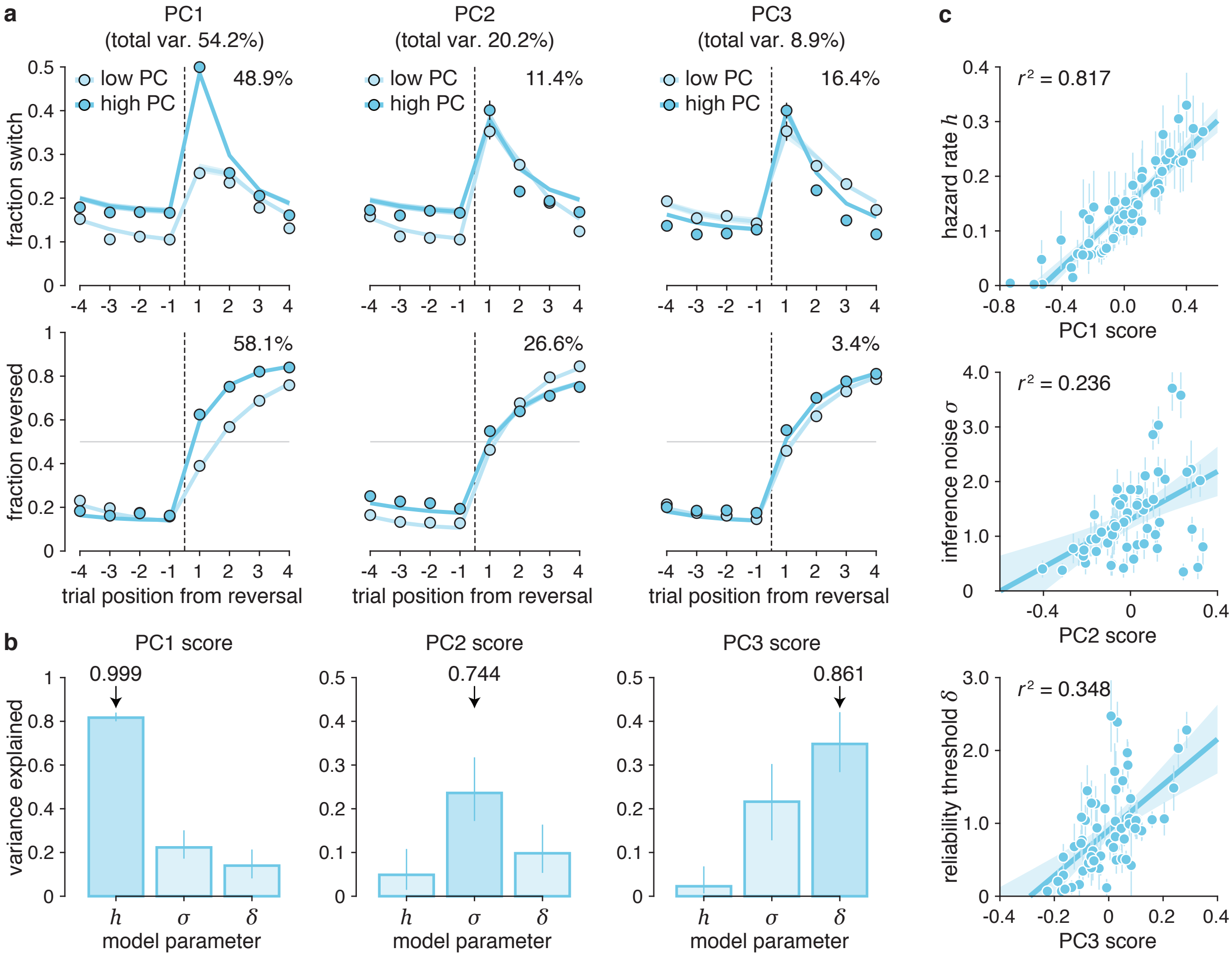


**Supplementary Figure 7 | Principal Component Analysis of interindividual variability** ($N = 60$). **(a)** Behavioral gradients associated with the first three PCs. Median-split on each principal component score for conditional inference model simulations (lines) and observed data (dots) and corresponding variance explained by each principal component. Error bars and shaded areas correspond to s.e.m. **(b)** Relations between each PC and model parameters. Each conditional inference best-fitting model parameter explains best the variance of one principal component score. The perceived hazard rate best explains the first PC score with exceedance $p_{\text{exc}} > 0.999$, the inference noise best explains the second PC score with exceedance $p_{\text{exc}} > 0.744$ and the reliability threshold best explains the third PC score with exceedance $p_{\text{exc}} > 0.861$. Bars correspond to $r^2$ and error bars to the interquartile ranges of each $r^2$ measure obtained through bootstrapping ($N = 10^4$). **(c)** Relation between PC scores and their associated parameters. Each principal component score significantly correlates with one conditional inference model parameter. Solid blue line correspond to the best-fitting regression line (in the least-squares sense) and shade area corresponds to the 95% confidence interval for predicted values. Parameters are presented as mean ± s.d. of posterior distributions.

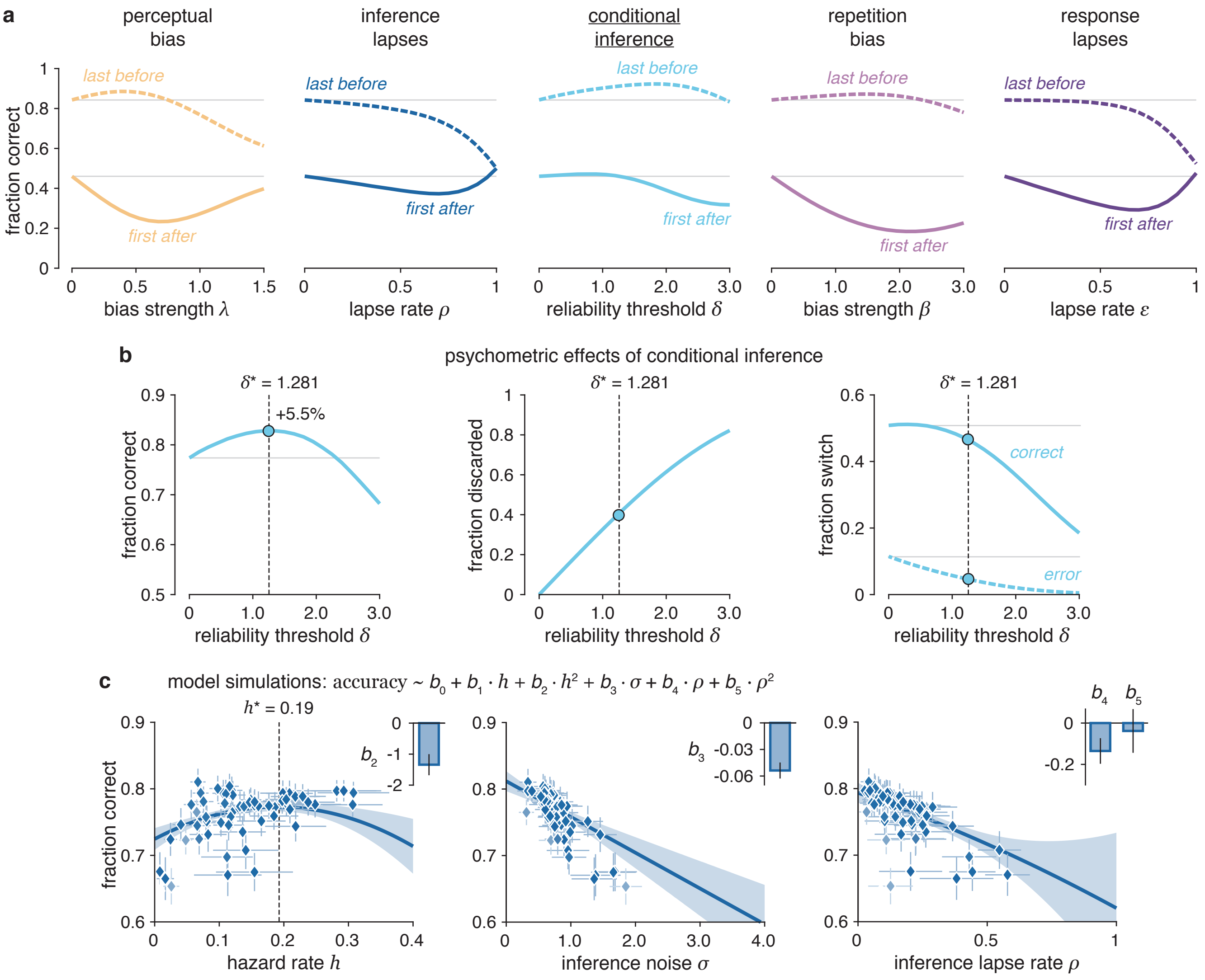


**Supplementary Figure 8** | **Conditional inference model effects. (a)** Simulated effects on response accuracy before and after a reversal. Accuracy computed on the trial just before (dotted lines) and just after (solid lines) a reversal depending on each stabilization parameter. All stabilization parameters reduce accuracy on the trial following a reversal except for the conditional inference model which maintains the accuracy stable. The asymptotic accuracy reached before reversal is increased by three stabilization parameters: bias strength, reliability threshold and bias strength with a stronger and systematic improvement provided by the conditional inference model. (b) Psychometric effects of conditional inference. Left: Overall accuracy is increased when introducing a conditional threshold by +5.5% for $\delta^* = 1.281$ (see also main Figure 9a). Middle: fraction of discarded evidence depending on the reliability threshold (41.3% for $\delta^* = 1.281$). Right: Fraction of switch leading to a correct response (solid line) and leading to an error (dotted line). **(c)** Predicted effects of best-fitting model parameters on inference lapses modeled decision accuracy (robust multiple regression: $r$-squared $= 0.888$, $p < 0.001$). For each best-fitting inference lapse model parameter, corresponding inference lapse model predicted accuracy (diamonds) and robust regression (solid blue lines). The lapse rate quadratic regressor $b_5$ is not significant ($p = 0.72$), therefore no inverted U-shaped relation to accuracy emerges. Solid blue line corresponds to the best-fitting robust regression line (in the weighted least-squares sense). Shaded areas correspond to the 95% confidence interval for predicted values. Parameters are presented as mean $\pm$ s.d. of posterior distributions and vertical error bars correspond to the accuracy s.d. of best-fitting model simulations. Top right insets correspond to estimated regression coefficients presented as mean $\pm$ s.e.m.

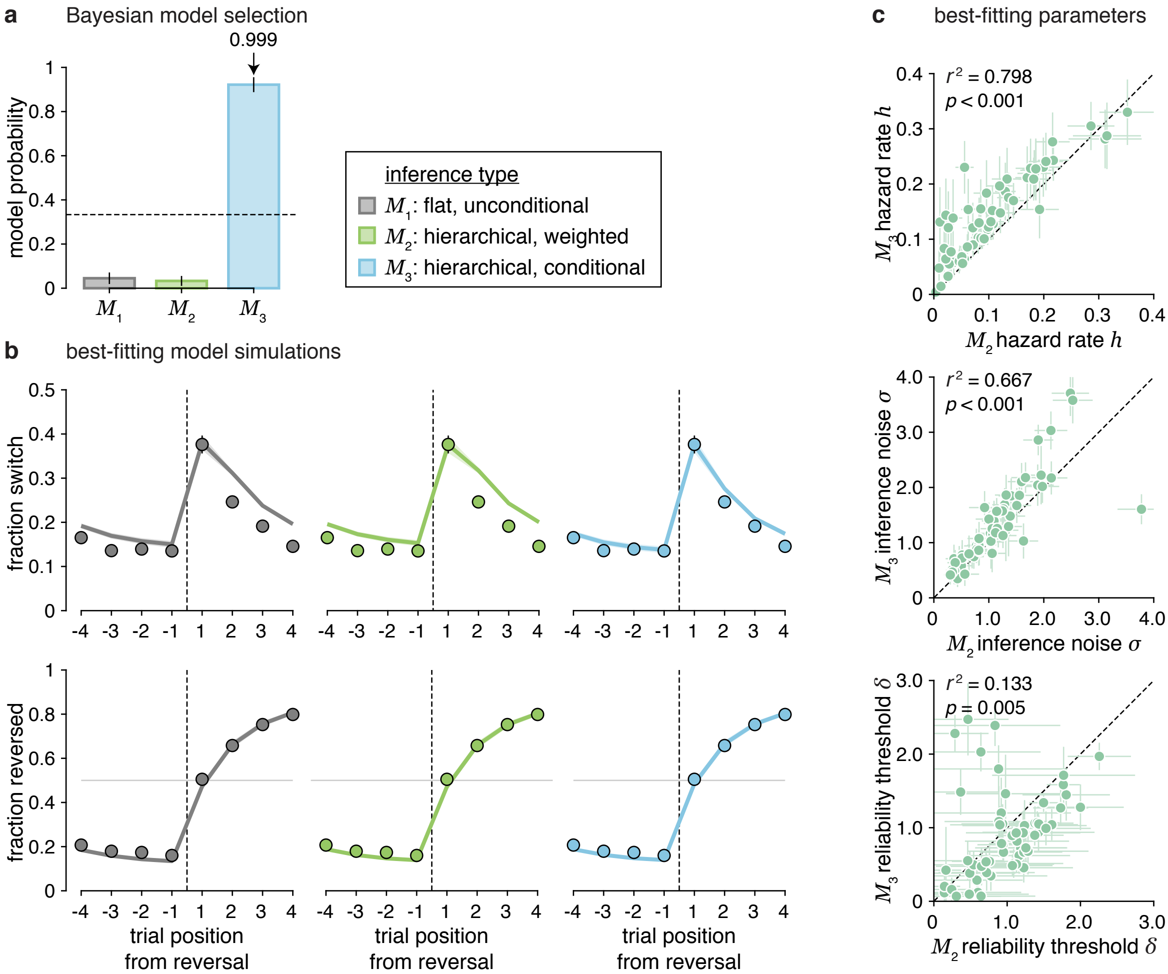


**Supplementary Figure 9 | Comparison between flat unconditional, hierarchical weighted and hierarchical conditional inference models ($N$ = 60). (a)** Estimated model probabilities for flat, unconditional inference, for hierarchical, weighted, and for hierarchical, conditional inference when fitted to human data from both experiments pooled, random-effects Bayesian model selection provides clear support for the hierarchical conditional inference model with exceedance probability $p_{\text{exc}} > 0.999$. Model probabilities are presented as mean and s.d. of the estimated Dirichlet distribution. Dashed lines correspond to the uniform distribution. **(b)** Simulations of flat unconditional, of hierarchical weighted and of hierarchical conditional inference model fitted to participants' (experiments pooled). Simulated response switch curves (top row) and response reversal curves (bottom row) based on the best-fitting parameters of each model. Solid colored lines correspond to the reversal behavior of each stabilized model whereas dots indicate human data (group-level average). If the flat unconditional model reproduces well participants' reversal behavior for both metrics, especially on the trial just following a reversal for the response switch curve – unlike most other stabilization models considered in this paper – it fails at reproducing participants' stabilized behavior on the following trials as well as the hierarchical conditional model. The hierarchical weighted inference model reproduces well participants' fraction reverse but fails at reproducing their stabilized behavior. Shaded areas and error bars correspond to s.e.m. **(c)** Best-fitting parameter correlations. Weighted and conditional hierarchical inference model have the same free parameters and their best-fitting values correlate positively. Parameters are presented as mean ± s.d. of posterior distributions. Dotted lines correspond to the identity line.

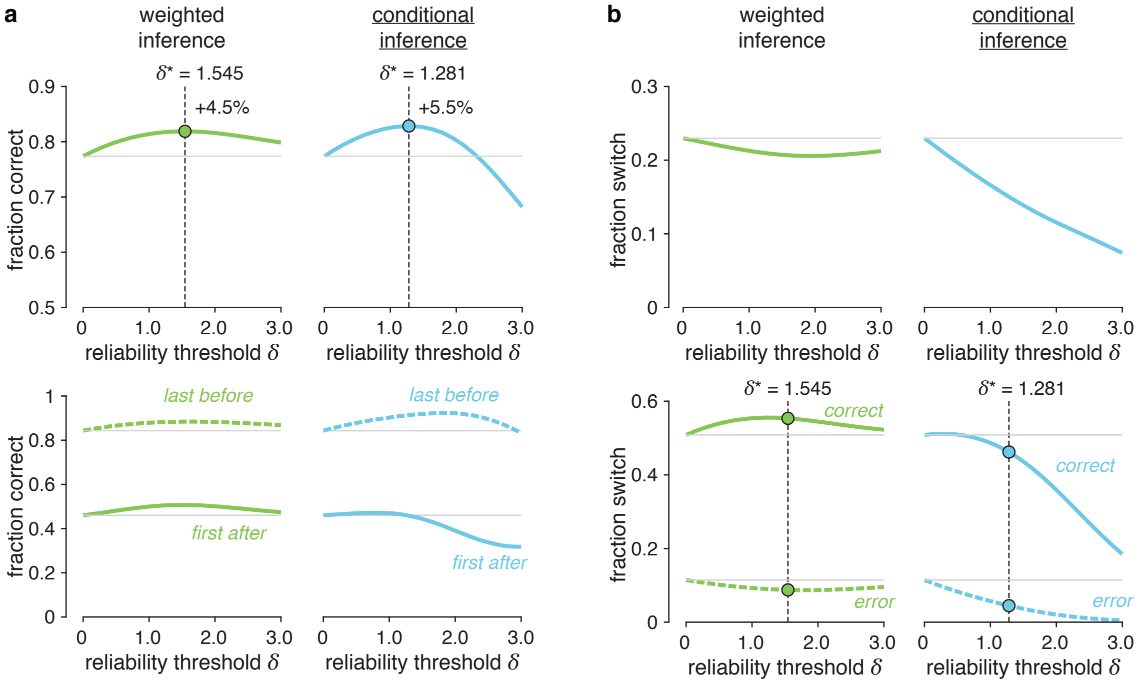


**Supplementary Figure 10 | Comparison between weighted inference and conditional inference model effects.** **(a)** Simulated effects of weighted and conditional inference models on decision accuracy. Top: Fraction of overall correct responses when increasing the reliability threshold. The accuracy is increased by +4.5% for $\delta^* = 1.55$ for the weighted inference model, a little less than for the conditional inference model increasing the accuracy by +5.5% for $\delta^* = 1.28$. Dots correspond to the best simulated accuracy. Down: Simulated effects on response accuracy before and after a reversal. Accuracy computed on the trial just before (dotted lines) and just after (solid lines) a reversal depending on the reliability threshold for weighted and conditional inference models. Both models increase the asymptotic accuracy reached before a reversal and the weighted inference model maintain the accuracy even more stable on the trial after a reversal than the conditional inference model. **(b)** Simulated effects of weighted and conditional inference models on switching behavior. Top: weighted inference barely reduces overall behavioral variability compared to the conditional inference model. Infinitely large values of the reliability threshold predict the same effect on the fraction switch as no reliability threshold at all. Bottom: Fraction of switch leading to a correct response (solid line) and leading to an error (dotted line) for both models.

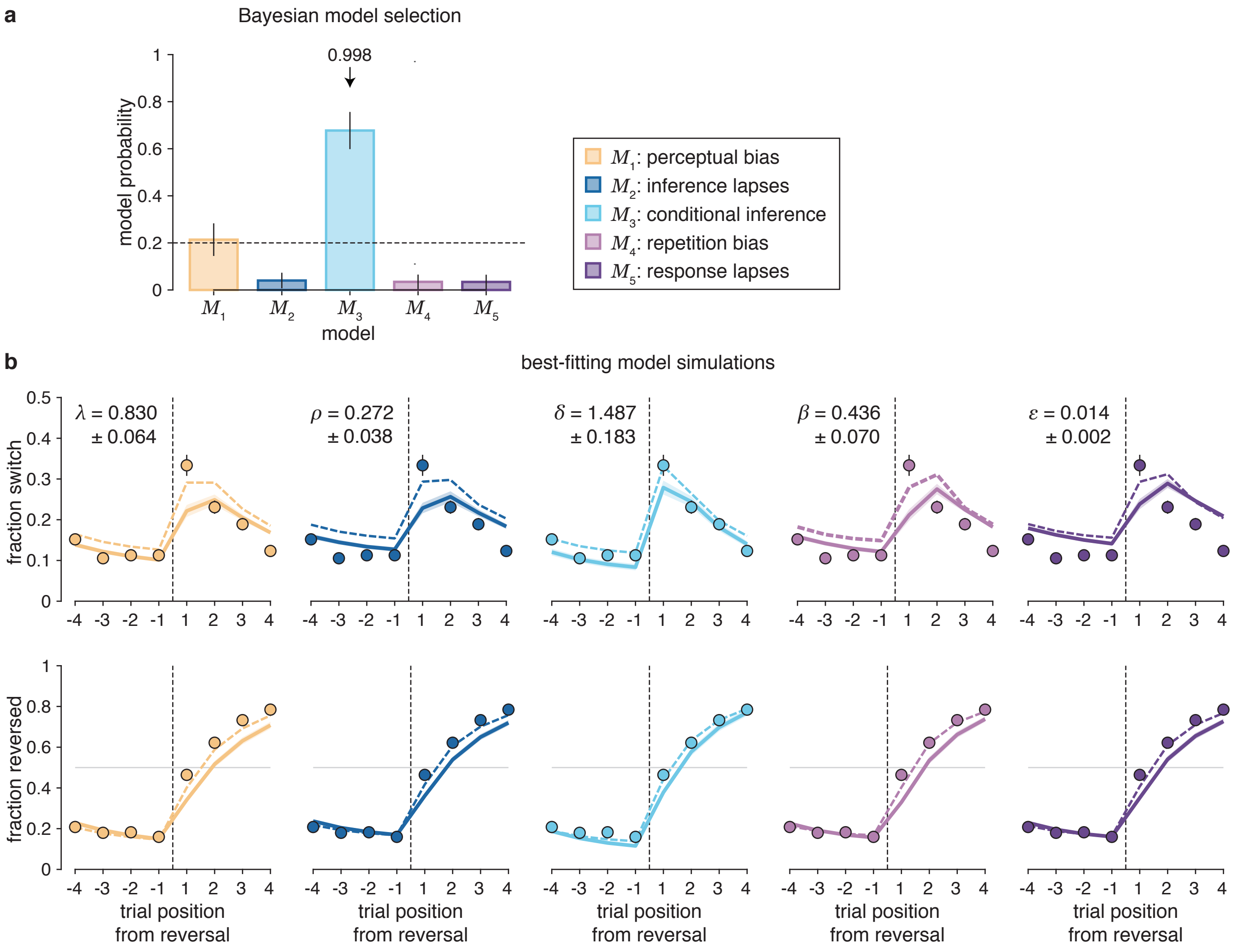


**Supplementary Figure 11 | Belief stabilization through conditional inference – fit to each choice** ($N = 30$) **(a)** Random-effects Bayesian model selection of the best-fitting strategy in each participants' choice – instead of focusing only on the two reversal metrics. Bars indicate the estimated model probabilities for the five candidate strategies for experiment 1. Again, the conditional inference model ($M_3$) outperforms other candidate strategies with exceedance $p_{\text{exc}} > 0.999$. Model probabilities are presented as mean and s.d. of the estimated Dirichlet distribution. Dashed line corresponds to chance probability. **(b)** Simulations of response stabilization strategies fitted to each participants' choice. Simulated response switch curves (top row) and response reversal curves (bottom row) based on the best-fitting parameters of each model. The mean best-fitting stabilizing parameter ± s.e.m. is indicated in the top-left corner for each model. Solid colored lines correspond to the reversal behavior of each stabilized model with parameters best-fitting each participant's choice, whereas dashed colored lines correspond to the reversal behavior of each stabilized model with parameters best-fitting these characteristic two metrics around reversal (same as main Figure 4). Dots indicate human data (group-level average). Again, the conditional inference model reproduces participants' reversal behavior better than any other models. Both curves are qualitatively less accurate when maximizing choice probability log-likelihood instead of these both metrics log-likelihoods. Shaded areas and error bars correspond to s.e.m.

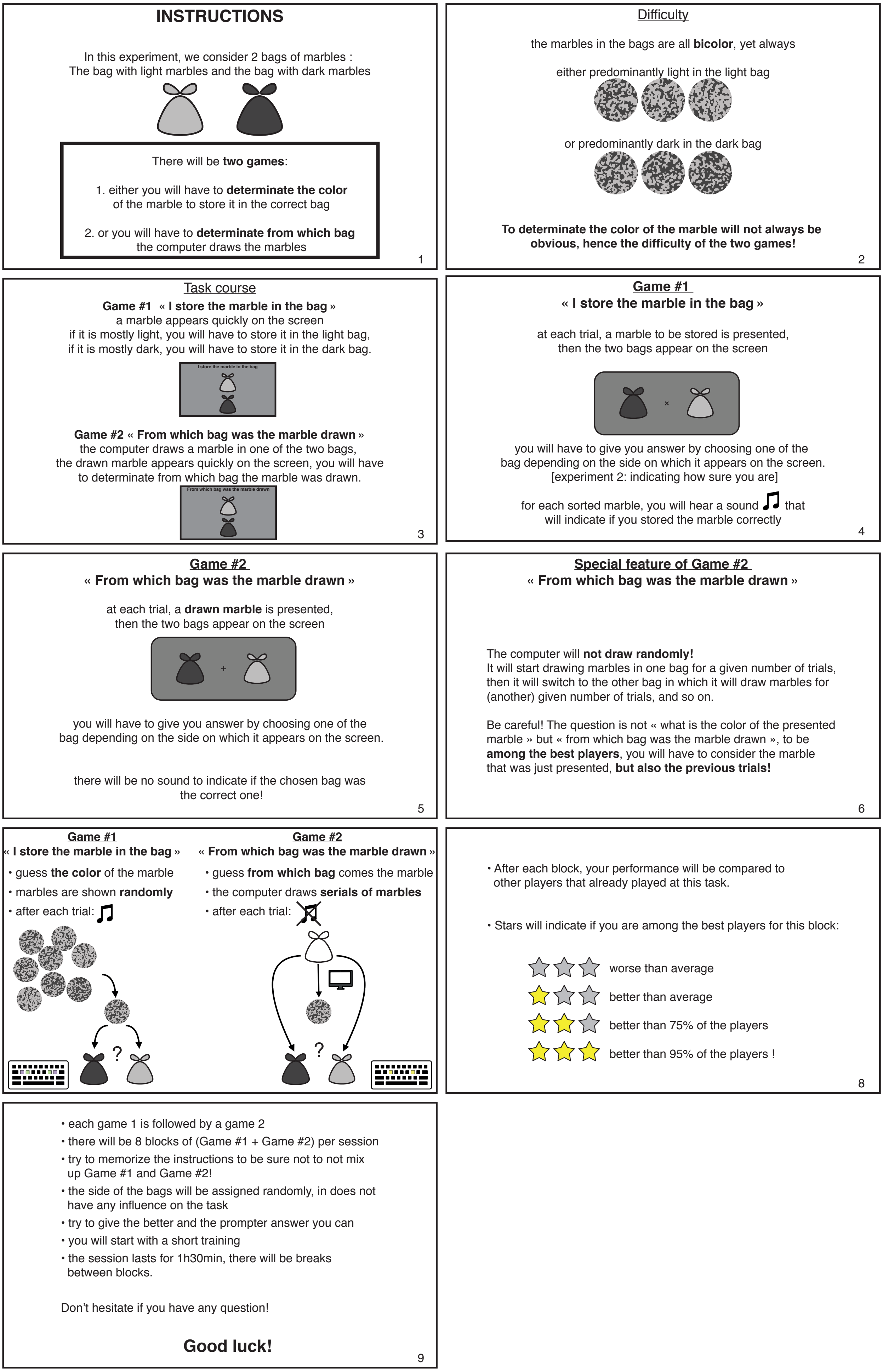


**Supplementary Figure 12 | Task instructions.**
